# Population-Scale Sequencing Data Enables Precise Estimates of Y-STR Mutation Rates

**DOI:** 10.1101/036590

**Authors:** Thomas Willems, Melissa Gymrek, G. David Poznik, Chris Tyler-Smith, The 1000 Genomes Project Chromosome Y Group, Yaniv Erlich

## Abstract

Short Tandem Repeats (STRs) are mutation-prone loci that span nearly 1% of the human genome. Previous studies have estimated the mutation rates of highly polymorphic STRs using capillary electrophoresis and pedigree-based designs. While this work has provided insights into the mutational dynamics of highly mutable STRs, the mutation rates of most others remain unknown. Here, we harnessed whole-genome sequencing data to estimate the mutation rates of Y-chromosome STRs (Y-STRs) with 2-6 base pair repeat units that are accessible to Illumina sequencing. We genotyped 4,500 Y-STRs using data from the 1000 Genomes Project and the Simons Genome Diversity Project. Next, we developed MUTEA, an algorithm that infers STR mutation rates from population-scale data using a high-resolution SNP-based phylogeny. After extensive intrinsic and extrinsic validations, we harnessed MUTEA to derive mutation rate estimates for 702 polymorphic STRs by tracing each locus over 222,000 meioses, resulting in the largest collection of Y-STR mutation rates to date. Using our estimates, we identified determinants of STR mutation rates and built a model to predict rates for STRs across the genome. These predictions indicate that the load of de novo STR mutations is at least 75 mutations per generation, rivaling the load of all other known variant types. Finally, we identified Y-STRs with potential applications in forensics and genetic genealogy, assessed the ability to differentiate between the Y-chromosomes of father-son pairs, and imputed Y-STR genotypes.

## Introduction

Mutations provide the fuel for evolutionary processes. The rates at which new mutations arise play a central role in a range of genetic applications, including dating phylogenetic events^1^, informing disease studies^2^, and evaluating forensic evidence.^3^ The advent of high-throughput sequencing has enabled genome-wide measurements of the number of de novo mutations using a broad range of strategies. A host of studies have evaluated the mutation rates of nearly every type of genetic variation, ranging from SNPs^4–7^ and short indels^8^ to large structural variations.^9^ These sequencing studies have concluded that approximately 50-100 de novo mutations arise each generation, most of which are point mutations. However, these studies have largely overlooked the contribution of short tandem repeats (STRs).

STRs are one of the most abundant types of repeats in the human genome. They consist of a repeating 2-6 base pair (bp) motif and span a median of 25bp. Approximately 700,000 STR loci exist in the human genome that in aggregate occupy ~1% of its total length. STR variations have been implicated in more than 30 hereditary disorders^10^, and emerging lines of evidence have highlighted their involvement in complex traits in both humans^11–13^ and model organisms.^14–16^ The repetitive nature of STRs causes error-prone DNA-polymerase replication events that can insert or delete copies of the repeat motif in subsequent generations, leading to markedly elevated mutation rates.^17; 18^

Previous studies estimated the rates and patterns of de novo STR mutations using capillary electrophoresis genotyping of specialized sets of markers, such as the Marshfield panel, the CODIS markers, or specific Y-chromosome STRs (Y-STRs). These studies have estimated that the average STR mutation rate per locus is 10^−3^ to 10^−4^ mutations per generation (mpg).^17; 19–22^ However, most STRs characterized in these studies were chosen for their relatively high levels of diversity in the population. As such, it is not clear whether their mutation rates and patterns reflect most STRs in the genome. Furthermore, as most previously studied STRs have tri- and tetranucleotide motifs, the field lacks robust mutation rate estimates for other motif lengths, specifically dinucleotides, the most prevalent type of STR. Finally, capillary electrophoresis has relatively low throughput, and most STRs were never genotyped in these studies, leaving the specific mutation rates of most STRs unknown.

The rapid advancement of next-generation sequencing technologies has provided the opportunity to genotype STRs beyond those on existing panels and to do so on a larger scale. Coupled with vast improvements in the depth, read length, and quality of whole-genome sequencing (WGS) datasets, algorithmic progress in STR genotyping tools has made it possible to robustly call these markers from high-throughput data.^23–25^ In our previous study, we found that 90% of the STRs in the genome are accessible to Illumina technology, and we showed that hemizygous STRs can be called with very high accuracy.^26^

Here, we leveraged population-scale high-throughput sequencing data to systematically estimate the mutation rates and analyze the mutational dynamics of STRs across the Y-chromosome. To gain power, we used two independent datasets, the 1000 Genomes Project^27^ and the Simons Genome Diversity Project (SGDP).^28^ The Y-chromosomes in these datasets confer rich genealogical information, enabling the analysis of complex STR mutation models without the need for familial information. To leverage this genealogical information, we developed an algorithm, Measuring Mutation Rates using Trees and Error Awareness (MUTEA), which infers the mutational dynamics along the Y-chromosome branches. After validating MUTEA via intrinsic and extrinsic tests, we scanned 4,500 Y-STRs and used the algorithm to infer the mutation rates of 702 polymorphic Y-STRs. To the best of our knowledge, this is the largest collection of Y-STR mutation rates to date. We show the value of this large collection of mutation rates by uncovering the sequence determinants of mutability, predicting the genetic load of de novo STR mutations across the genome, and exploring a series of forensic applications.

## Materials and Methods

### Sequencing Datasets

We analyzed 179 male samples in the SGDP cohort from widely dispersed populations across Africa, Asia and the Americas. The SGDP samples were sequenced to over 30× coverage using a PCR-free library preparation protocol and 100bp paired-end Illumina reads. As our previous results demonstrate that this protocol substantially reduces the rate of PCR stutter at STR loci^29^, the SGDP cohort provides a high-quality dataset for calling Y-STRs. We also analyzed 1,244 unrelated male samples from phase 3 of the 1000 Genomes Project. These samples are from 26 globally diverse populations and were sequenced to an average autosomal coverage of 7**x** using 75-100 bp paired-end Illumina reads.

### Y-SNP Phylogeny

To construct the SGDP Y-chromosome haplotype tree, we downloaded VCF files containing the Y-SNP calls generated by the SGDP analysis group. As many of these SNPs lie in pseudoautosomal regions or regions with low mappability, we applied a series of filters to reduce the frequency of genotyping errors. We first removed loci where more than 10% of individuals were heterozygous using VCFtools.^30^ For the remaining SNPs, we removed individual SNP calls that were heterozygous, had fewer than 7 supporting reads, or had more than 10% of reads supporting an uncalled allele. Lastly, we discarded SNP loci if fewer than 150 samples met these criteria or if more than 10% of reads had zero mapping quality. Overall, we obtained nearly 39,000 high-quality polymorphic SNPs.

We then used the high-quality SNPs to build the Y-chromosome phylogenetic tree using RAxML^31^ and the options –`m ASC_GTRGAMMA –f d –asc-corr lewis`. The SGDP samples included 3 representatives of haplogroup A1b1 and no members of the more basal clades (A00, A0, and A1a), so we used Dendroscope^32^ to root the phylogeny along the branch marked by the M42 and M94 mutations, markers associated with the split between A1b1 and megahaplogroup BT. For the 1000 Genomes phase 3 dataset, we used a RAxML-generated phylogeny that was built by the 1000Y analysis group.^33^

Although the maximum-likelihood phylogeny generated for each dataset has numerical branch lengths, these lengths are not scaled in units of generations as required by our method. We therefore tested two scaling approaches. First, we selected the factor that most closely equated the total number of generations in each phylogeny to the corresponding value based on published Y-SNP mutation rates. To do so, we used a recently published Y-SNP mutation rate of 3×10^−8^ mutations per base per generation^34; 35^ and the numbers of called SNPs and called sites in each SNP dataset. As an alternative method, we scaled the trees using mutation rate estimates for 15 loci in the Y-chromosome Haplotype Reference Database (YHRD), a large compendium of individual Y-STR mutational studies (individually cited therein).^36^ We chose to calibrate using these loci because their mutation rate estimates are each based on more than 7,000 father-son pairs per locus and should therefore be relatively precise. For the 1000 Genomes data, we used the available PowerPlex capillary data for each locus, assumed error-free genotypes, scaled the phylogeny using a range of factors, and estimated the set of mutation rates for each scaling factor using MUTEA (see below). The choice of scaling factor had essentially no effect on the correlation with the YHRD estimates, resulting in an *R*^*2*^ of 0.89 across all tested factors (**Figure S1**). However, the total squared error between the estimates was minimized for a factor of ~2,800, which we therefore selected as the optimal scaling. For the SGDP data, we performed an analogous analysis using HipSTR genotypes (see below) for 9 of these 15 loci, again resulting in a uniform *R*^*2*^ of 0.91 and an optimal scaling factor of ~3,200 (**Figure S1**).

The resulting scaling factors were remarkably concordant between the methods, with the factors determined by the Y-SNP method ~25% greater. However, to maximize the concordance with pedigree estimates, we used the second method. After scaling the branches, we found that the approximate total lengths of the SGDP and 1000 Genomes phylogenies are 60,000 and 160,000 meioses, respectively.

### Defining and Identifying Y-STRs

To identify Y-STRs, we used a quantitative procedure developed in our previous work.^26^ Briefly, this procedure uses Tandem Repeats Finder (TRF) to score each genomic sequence according to its purity, length, and nucleotide composition.^37^ It then uses extensive simulations of random nucleotide sequences to determine a scoring threshold that distinguishes random DNA from DNA that is truly repetitive, selecting regions with scores above this threshold as STRs. Our previous results suggest that this approach has less than a 1.4% probability of omitting a polymorphic STR and has a false positive rate of approximately 1%.

We applied this procedure to the Y-chromosome sequence of the hg19 reference genome. As TRF occasionally identifies regions that overlap, we ensured that every locus has a unique STR annotation using the following steps: (1) We merged two STR regions if the higher scoring one contained 85% of the bases in the union of the regions (2) Overlapping entries that failed this criterion but which had the same period were also merged. For example, adjacent [GATA]10 and [TACA]8 entries were merged into one STR (3) Since we intended to use sequencing alignments relative to either hg19 or GRCh38 coordinates, we removed hg19 STR regions that failed to liftOver^38^ to the GRCh38 assembly or were lifted from the Y-chromosome to the X-chromosome.

We also added coordinates for Y-STR loci whose mutation rates have been characterized in prior studies.^21; 39^ For these markers, we used the published set of primer sequences and the isPCR tool^38^ to map the primers to hg19 coordinates. We then ran TRF on each region and pinpointed the coordinates using the published repeat structure. Lastly, we applied TRF to additional regions previously published as part of comprehensive Y-STR maps to obtain coordinates for labeled markers whose mutation rates have not been characterized.^40^ In total, we added 261 annotated Y-STRs, ~190 of which have mutation rate estimates from prior studies. The complete Y-STR reference is available for download in both hg19 and GRCh38 coordinates (**Web resources**).

### Y-STR Call Set and its Accuracy

We downloaded BWA-MEM^41^ alignments for the SGDP samples from the project website and extracted and merged the Y-chromosome alignments into a single BAM file using SAMtools.^41^ STR genotypes were then generated using HipSTR, an improved version of lobSTR, an STR caller for Illumina data we developed in our previous studies.^23^

HipSTR provides additional capabilities over lobSTR by using a specialized hidden Markov model (HMM) to account for PCR stutter artifacts. Briefly, to genotype an STR, HipSTR creates a list of candidate alleles from the alignments observed in the population. For each sample, it then realigns every read to each putative allele using the HMM, selects the allele with the highest total likelihood as the genotype, and returns each read’s alignment relative to this genotype. This haplotype-based approach produces highly accurate STR genotypes and eliminates many read misalignments that occur if reads are aligned individually or are only aligned to the reference genome. We used HipSTR to genotype each STR region in the Y-STR reference described above using the merged BAMs and the following options: `‐‐min-reads 25 ‐‐haploid-chrs chrY ‐‐hide-allreads`. Similarly, we downloaded BWA-MEM alignments from the 1000 Genomes phase 3 data release. As these alignments were relative to the GRCh38 assembly, we ran HipSTR using the corresponding GRCh38 STR regions and the options `‐‐min-reads 100 ‐‐haploid-chrs chrY ‐‐hide-allreads`.

We employed several strategies to enhance the quality of the SGDP STR call set: (1) To avoid errors introduced by neighboring repeats, we omitted genotyped loci that overlapped one another or multiple STR regions (2) We discarded loci if more than 5% of samples’ genotypes had a noninteger number of repeats, such as a three base pair expansion in an STR with a tetranucleotide motif. These types of events occur quite rarely and usually reflect genotyping errors rather than genuine STR polymorphisms^23^ (3) We removed Y-STRs sites that were called in at least 2 SGDP females, as they are likely to have high X-chromosome or autosome homology (4) We omitted sites if more than 15% of reads had a stutter artifact or more than 7.5% of reads had in indel in the sequence flanking the STR. These HipSTR-reported statistics typically indicate that the locus is not well captured by HipSTR’s genotyping model and may arise if duplicated sites are mapping to the same reference genome location (5) For the remaining loci, we discarded unreliable calls on a per-sample basis if more than 10% of an individual’s reads had an indel in the flank sequence (6) Finally, we removed loci in which fewer than 100 samples had genotype posteriors greater than 66%, as these loci had too few samples for accurate inference.

To filter the 1000 Genomes call set, we first removed loci that did not pass the SGDP dataset filters. We then applied a set of filters identical to those described above except that we only removed loci with more than 15 genotyped females and did not apply a stutter frequency cutoff. These alterations account for the 1000 Genomes dataset’s larger sample size and use of PCR amplification during library preparation.

Importantly, we found that both the SGDP and 1000 Genomes HipSTR call sets had high quality. We compared our STR genotypes to capillary electrophoresis datasets available for the same samples. For the SGDP, we observed a 99.7% concordance rate when comparing the HipSTR and capillary results for 3,300 calls at 48 Y-STRs.^42^ For the 1000 Genomes, a comparison of 4,050 calls at 15 loci in the PowerPlex Y23 panel resulted in a 97.5% concordance rate.^43^

### Measuring Mutation Rates Using Trees and Error Awareness (MUTEA): Theory

Previously developed methods estimate STR mutation rates from population data by comparing the mean squared difference in allele lengths between samples to the time to the most recent common ancestor (TMRCA).^44; 45^ However, these methods generally assume simple mutation models, can be sensitive to haplogroup size fluctuations^46^ and require exact error-free genotypes. We therefore sought to develop an algorithm that can address these issues by leveraging detailed Y-SNP phylogenies.

**Figure 1** outlines the steps underlying MUTEA. Under a naïve setting without genotyping error, MUTEA uses Felsenstein’s pruning algorithm^47^ and numerical optimization to evaluate and improve the likelihood of a mutation model until convergence. However, due to the error-prone and low-coverage nature of WGS-based STR call sets, using these genotypes would result in vastly inflated mutation rate estimates. To avoid these biases, MUTEA learns a locus-specific error model and uses this error model to compute genotype posteriors. It then uses these posteriors rather than fixed genotypes during the mutation model optimization process to obtain robust estimates. In addition, MUTEA uses a flexible computational framework for STR mutations that includes length constraints and allows for multi-step mutations. We describe each step below.

**Figure 1:**
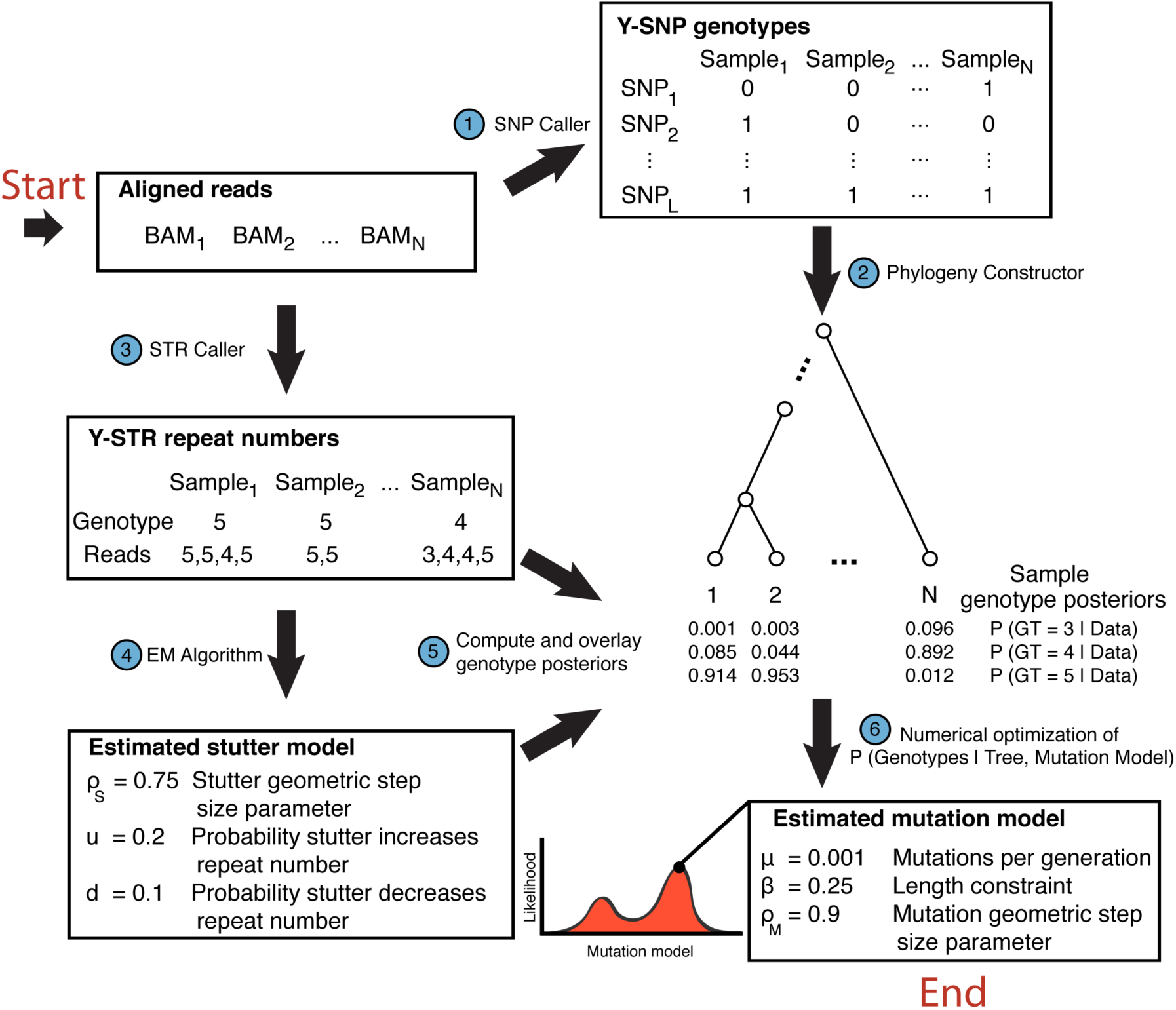
Y-STR mutation rate estimation method. Schematic of our procedure to estimate Y-STR mutation rates. The method first genotypes Y-SNPs (step 1) and uses these calls to build a single Y-SNP phylogeny (step 2). This phylogeny provides the evolutionary context required to infer Y-STR mutational dynamics, with samples in the cohort occupying the leaves of the tree and all other nodes representing unobserved ancestors. Steps 3-6 are then run on each Y-STR individually. After using an STR genotyping tool to determine each sample’s maximum-likelihood genotype and the number of repeats in each read (step 3), an EM-algorithm analyzes all of these repeat counts to learn a stutter model (step 4). In combination with the read-level repeat counts, this model is used to compute each sample’s genotype posteriors (step 5). After randomly initializing a mutation model, Felsenstein’s pruning algorithm and numerical optimization are used to repeatedly evaluate and improve the likelihood of the model until convergence. The mutation rate in the resulting model provides the maximum-likelihood estimate.

### Mutation Model Likelihood

We used Felsenstein’s pruning algorithm to evaluate the likelihood of an STR mutation model. Let *M* denote the STR mutation model, *D* denote the dataset containing STR genotype likelihoods, and *T* denote the Y-chromosome phylogeny rooted at node *R*. The likelihood of the data is:

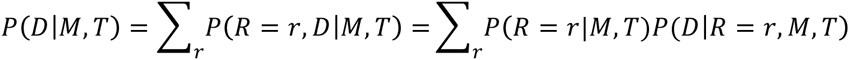

Let *D*_*N*_*i*__ denote the genotype likelihoods of all nodes that are in the subtree rooted at node *N*_*i*_. If node *N*_*i*_ has genotype *g*, the conditional probability of the data in its subtree is given by:

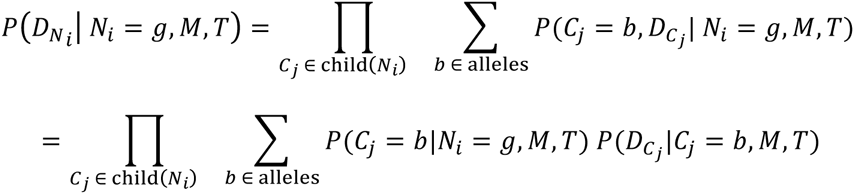

While descending the phylogeny, this recursive relation applies until a node with no children is encountered. These leaf nodes represent sequenced individuals and the conditional probability of the data is given by the individuals’ genotype likelihoods. Therefore, the likelihood of a mutation model can be calculated using a post-order tree traversal. First, the algorithm computes the genotype likelihoods at each leaf node. It then progresses to each internal node and calculates the conditional probability of the data for each potential genotype after computing its descendants’ probabilities. Finally, upon reaching the root node, the total data likelihood is computed using the root node’s conditional probabilities and a uniform prior for the root node’s genotype.

In practice, we compute the total log-likelihood to avoid numerical underflow issues. Because normalizing the genotype likelihoods of each sample does not affect the relative model likelihoods, we calculated genotype posteriors using a uniform prior and used them throughout our analysis.

### STR Mutation Model

To model STR mutations, we used a generalized stepwise mutation model with a length constraint. Each mutation model *M* is characterized by three parameters: a per-generation mutation rate *μ*, a geometric step size distribution with parameter *ρ*_*M*_ and *β*, a spring-like length constraint that causes alleles to mutate back towards the central allele. In this framework, the central allele is assigned a value of zero, and nonzero allele values indicate the number of repeats from this reference point. Given a starting allele *a*_*t*_ observed at time *t*, the probability of observing a particular allele *k* the following generation is:

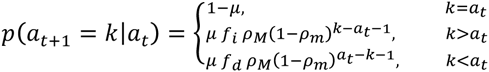

where the fraction of mutations increasing and decreasing the size of the STR are 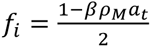 and *f*_*d*_ = 1 – *f*_*i*_; *f*_*i*_ values greater than one or less than zero were clipped and set to one and zero, respectively. These two model features act as spring-like length constraints that attract alleles back towards the central allele. To avoid biologically implausible models, we constrained *β* to have non-negative values, where *β* = 0 reduces to a traditional generalized stepwise mutation model and increasingly positive values of *β* model STRs with stronger tendencies to mutate back towards the central allele. Values of *ρ*_*M*_ close to one primarily restrict models to single-step mutations, while smaller values of this parameter enable frequent multistep mutations.

### Computing STR Genotype Likelihoods

To calculate the likelihood of the data *D* observed in the leaf nodes, we needed to account for STR genotyping errors. These errors are mainly caused by PCR stutter artifacts that insert or delete STR repeat units in the observed sequencing reads. We therefore developed a method to learn each STR’s distinctive stutter noise profile.

Let *Θ*_*x*_ denote the stutter model for STR locus *x*.*Θ*_*x*_ is parameterized by the frequency of each STR allele (*F*_*i*_), the probability that stutter adds (*u*) or removes (*d*) repeats from the true allele in an observed read, and a geometric distribution with parameter *ρ*_*s*_ that controls the size of the stutter-induced changes. Given a stutter model and a set of observed reads (*R*), the posterior probability of each individual’s haploid genotype is:

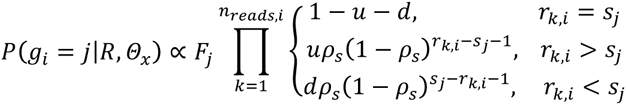

where *g*_*i*_ denotes the genotype of the *i*^*th*^ individual, *n*_*reads,i*_ denotes the number of reads for the *i*^*th*^ individual, *r*_*k,i*_ denotes the number of repeats observed in the *k*^*th*^ read for the *i*^*th*^ individual, and *s*_*j*_ denotes the number of repeats in the *j*^*th*^ allele. Analogous to the step size parameter in the mutation model, small values of *ρ*_*s*_ allow for frequent multistep stutter artifacts while values near one restrict artifacts to single step changes.

We implemented an expectation-maximization (EM) framework to learn these model parameters.^48^ The E-step computes the genotype posteriors for every individual given the observed reads and the current stutter model parameters. The M-step then uses these posterior probabilities to update the stutter model parameters as follows:

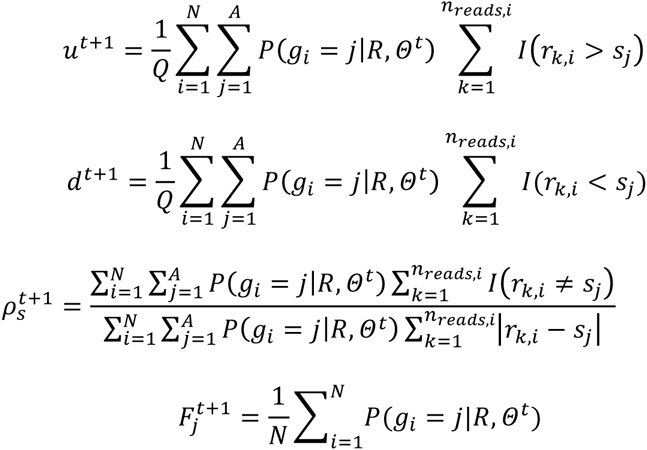

Here, *N* denotes the number of samples, *A* denotes the number of putative alleles, *Q* denotes the number of sequencing reads and *I* is the indicator function. As *ρ*_*S*_ is the parameter of a geometric step size distribution, the M-step updates its value using the inverse of the mean weighted step size for reads with non-zero stutter.

Locally misaligned reads can also introduce genotyping errors if they cause a miscalculation in a read’s repeat length. However, these errors introduce artifacts that are relatively similar to those caused by PCR stutter. As a result, the EM procedure learns stutter models that correct for the combined frequencies of PCR stutter and misalignment, resulting in robust genotype posteriors for downstream analyses.

### MUTEA Computation

Given genotype likelihoods for an STR of interest, we used a maximum-likelihood approach to estimate the underlying mutation model. Our approach first estimates the central allele of the mutation model by computing the median observed STR length and then normalizes all genotypes relative to this reference point. Next, it randomly selects mutation model parameters *μ*, *β*, and *ρ*_*M*_ subject to the constraint that they lie within the ranges of 10^−5^ to 0.05, 0 to 0.75 and 0.5 to 1.0, respectively. Using these bounds, the Nelder-Mead optimization algorithm^49^, and the outlined method for computing each model’s likelihood, we iteratively update the mutation model parameters until the likelihood converges. After repeating this procedure using three different random initializations to increase the probability of discovering a global optimum, our algorithm selects the optimized set of parameters with the greatest total likelihood.

For each STR in the SGDP and 1000 Genomes call sets that passed the requisite quality control filters, we first used the EM algorithm to learn a PCR stutter model. To run this algorithm, we obtained the size of the STR observed in each read from the MALLREADS VCF field. HipSTR uses this field to report the maximum-likelihood STR size observed in each read that spans its sample’s most probable haplotype. In conjunction with a uniform prior, the learned stutter model was then used to compute the genotype posteriors for each sample with a HipSTR quality score greater than 0.66. Samples with quality scores below this threshold were omitted because the genotype uncertainty can result in erroneous reported read sizes. Finally, together with the optimization procedure and the appropriate scaled Y-SNP phylogeny, we used these genotype posteriors to obtain a point estimate of the STR’s mutation rate.

## Results

### Verifying MUTEA using Simulations

We validated MUTEA’s inferences by running the algorithm on simulated data from a wide range of Y-STR mutation models (**Appendix A**). We tested mutation rates (*μ*) from 10^−5^ to 10^−2^ mpg, a range that encompasses most known polymorphic Y-STRs. We also varied the distribution of step-sizes for each STR mutation from a single step (*ρ*_*M*_ = 1) to a wide range of mutation steps (*ρ*_*M*_ = 0.75) and added various spring-like length constraints that ranged from no constraint (*β* = 0) to a strong attractor towards the central allele (*β* = 0.5).

MUTEA obtained unbiased estimates of the simulated mutation rate for nearly all scenarios (**Figure S2)**. We only observed a slight upward bias in the estimates for the slowest simulated mutation rate (*μ* = 10^−5^) due to the lower bound imposed during numerical optimization. In contrast, mutation rates estimated using simpler mutation models limited to single-step mutations or no length constraints were far more biased in these scenarios (**Figure S3**). MUTEA’s inferences were also robust to the presence of simulated PCR stutter noise. After forward simulating STRs, we simulated reads for each genotype and distorted their repeat numbers using various PCR stutter models (**Appendix B**). We then input these repeat counts into MUTEA instead of the STR genotypes. Although MUTEA was completely blind to the selected stutter parameters, it reported unbiased estimates of the Y-STR mutation rates, step sizes, and stutter models for nearly all scenarios (**Figure 2, Figures S4–6**), with just a slight bias for the lowest simulated mutation rate, as was the case for the exact genotypes scenario described above. As a negative control, we again ran MUTEA on the stutter-affected reads but without employing the EM stutter correction method. With this procedure, posteriors based on the fraction of reads supporting each genotype resulted in marked biases, particularly for low mutation rates, demonstrating the importance of correctly accounting for stutter artifacts in these settings (**Figure 2, Figures S5–6**).

**Figure 2:**
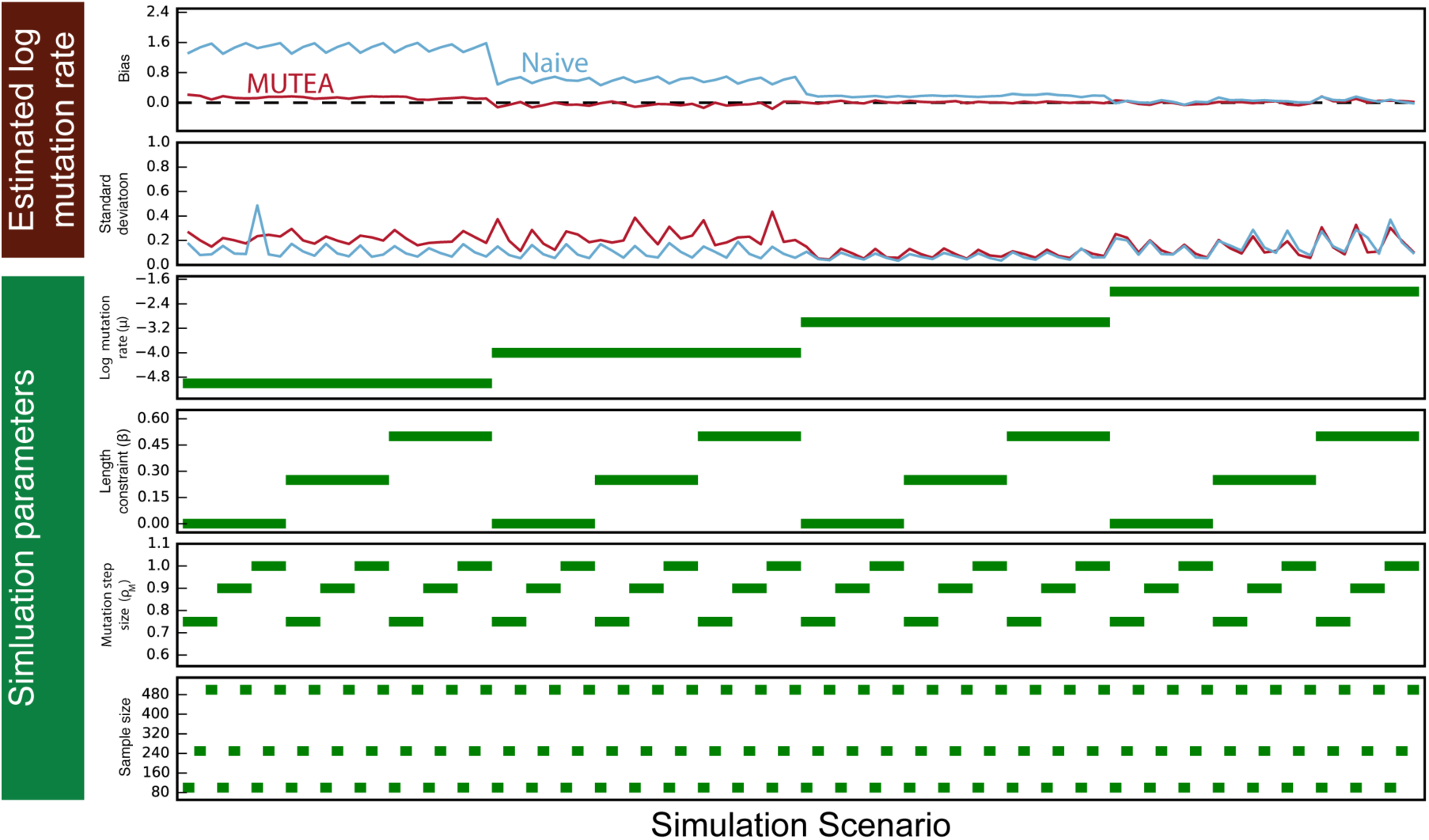
Validating MUTEA using simulations. STR sequencing reads with PCR stutter noise were simulated for a variety of sample sizes and mutation models (simulation parameters panels). Applying MUTEA (red line) to these reads led to relatively unbiased mutation rate estimates (upper panel) with small standard deviations (second panel). As a negative control, we also applied a naïve approach to correct for stutter noise (blue line). This approach computed genotype posteriors using the fraction of supporting reads, resulting in markedly biased mutation rate estimates.

### MUTEA Estimates are Internally and Externally Consistent

Encouraged by the robustness of our approach, we turned to analyze real Y-STR data from the SGDP and the 1000 Genomes Y-STR call sets. In total, we examined ~4,500 STR loci, 702 of which displayed length polymorphisms in both datasets, with the rest nearly fixed. We ran MUTEA on each of these polymorphic STRs to estimate its mutation rate (*μ*), expected step size (*ρ*_*M*_), and stutter parameters (*u*, *d*, *ρ*_*s*_) in both datasets (**Table S1**).

The MUTEA mutation rate estimates were largely consistent between the datasets (**Figure 3**). We obtained an *R*^*2*^ of 0.92 when comparing the log mutation rate estimates from the 1000 Genomes and SGDP datasets for the 702 polymorphic markers. Importantly, this high concordance was achieved despite substantial differences between the analyzed populations, sample sizes, and sequencing data quality. The 1000 Genomes data should have higher rates of stutter than the SGDP data due to the PCR amplification used in the sequencing library preparation.

**Figure 3:**
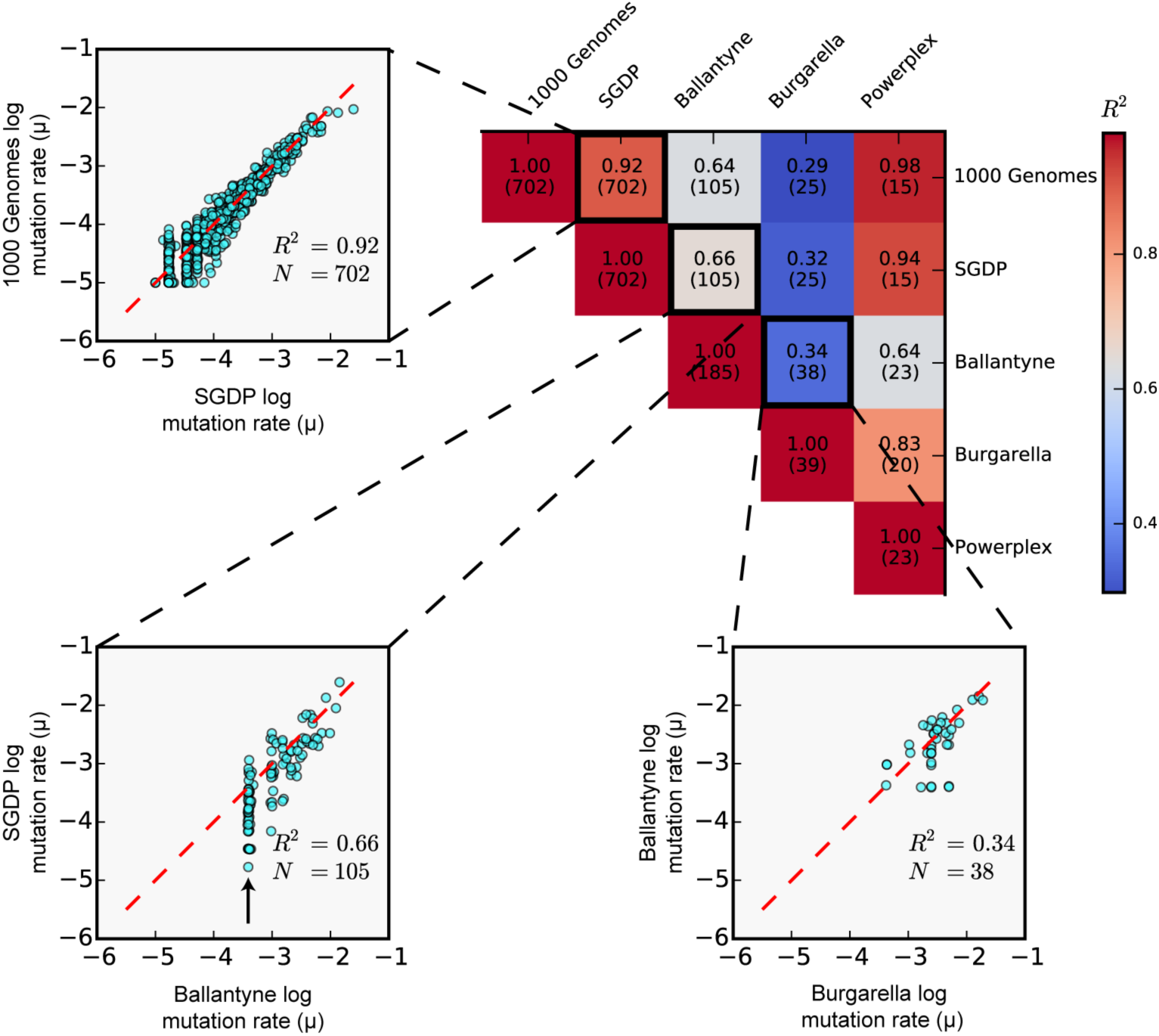
Concordance of mutation rate estimates across datasets. The heat map in the upper right corner presents the correlation of log mutation rates obtained from two father-son capillarY-based studies (“Ballantyne” and “Burgarella”) with those obtained in this study using the 1000 Genomes WGS data (“1000 Genomes”), the Simons Genome WGS data (“SGDP”) and the capillary data available for samples in the 1000 Genomes (“Powerplex”). Each cell indicates the number of markers involved in the comparison and the resulting *R*^*2*^. Representative scatterplots for three of these comparisons depict the pair of estimates for each marker (cyan) and the *x* = *y* line (red). The black arrow in the SGDP vs. Ballantyne comparison shows the effective lower limit of the Ballantyne et al. mutation rate estimates.

Consistent with this expectation, MUTEA learned higher stutter probabilities in the 1000 Genomes data, as compared to the SGDP data, for most loci (**Figure S7, left panels**). Nonetheless, the mutation rate estimates were highly concordant. In addition, we found that despite differences in the overall probability of stutter, the downward and upward stutter rates were highly correlated between the two datasets (*R*^*2*^ = 0.88 and *R*^*2*^ = 0.68 on the log scale, respectively), reflecting the algorithm’s ability to capture each locus’ distinctive error profile (**Figure S7, right panels)**.

Genotyping technology played only a small role in explaining the estimate concordance between the two datasets. We re-ran MUTEA on the 1000 Genomes Y-tree using capillary genotypes for 15 Y-STR loci that were available for the same samples (**Figure 3**). Comparing the resulting log mutation rate estimates to those obtained using sequencing-generated genotypes, we obtained an *R*^*2*^ of 0.98. These comparisons demonstrate that our method obtains robust locus-specific mutation rate estimates while accounting for varying degrees of PCR stutter artifacts and alignment and genotyping errors. Furthermore, the inter-dataset concordance suggests that there are either very few errors in the phylogenies or that these errors have little impact on the resulting mutation rate estimates.

We also validated our mutation rate estimates by comparing them to results from previous studies that used pedigree-based designs and capillary electrophoresis for genotyping. In these studies, Burgarella et al.^39^ and Ballantyne et al.^21^ estimated Y-STR mutation rates for specialized panels of Y-STRs by examining approximately 500 and 2,000 father-son duos per Y-STR, respectively. We observed only a moderate replicability between the reported mutation rates from these two prior studies (*R*^*2*^ of 0.34, **Figure 3**). This low correlation presumably stems from the very small number of transmissions used by Burgarella et al. On the other hand, we observed an *R*^*2*^ of ~0.65 when we compared either the SGDP or the 1000 Genomes estimates to those from Ballantyne et al., despite considerably different methodological approaches (**Figure 3**). One limitation of this comparison is that Ballantyne et al. could not report precise mutation rates for slowly mutating Y-STRs due to the number of meioses events examined in their study. As a result, their estimates were effectively restricted to a lower bound of *μ*=10^−35^ mpg (**Figure 3, inset**). In contrast, our deep phylogeny enabled us to accurately estimate much lower rates, highlighting the advantage of analyzing population data, rather than father-son pairs, for slowly mutating STRs. Comparing our estimates to those from Burgarella et al. resulted in an *R*^*2*^ of ~0.3, but restricting this evaluation to the subset of loci they characterized using more than 5000 father-son duos resulted in a substantially higher *R*^*2*^ of 0.87 (**Figure S8**). These results demonstrate that our estimates are concordant with prior father-son based results, provided that the latter were generated using sufficiently many pairs.

### Characteristics and Determinants of Y-STR Mutations

Next, we analyzed the STR mutation patterns. To obtain a single mutation rate estimate for each Y-STR, we averaged the estimates from the SGDP and 1000 Genomes datasets. We found that the distribution of Y-STR mutation rates has a substantial right tail, with most STRs mutating at very slow rates and only a few loci mutating at high rates (**Figure 4**). On average, a polymorphic Y-STR mutates at a rate of 3.8×10^−4^ mpg and has a median mutation rate of 8.7×10^−5^ mpg. The average Y-STR mutation rate is an order of magnitude lower than previous estimates from panel-based studies. This difference cannot be explained by our phylogenetic measurement procedure since inspection of the same markers yielded relatively concordant numbers. Instead, it likely stems from the ascertainment strategy of STR panels, which select highly diverse loci that do not reflect the mutation rates of most STRs. One caveat in this analysis is that very long Y-STR markers were not accessible to Illumina reads. These loci might affect the calculated average mutation rate and, to a smaller extent, the median mutation rate. Consistent with these explanations, our mutation rate estimates for previously characterized loci were upwardly enriched relative to our estimates for all markers (**Figure 4**).

**Figure 4:**
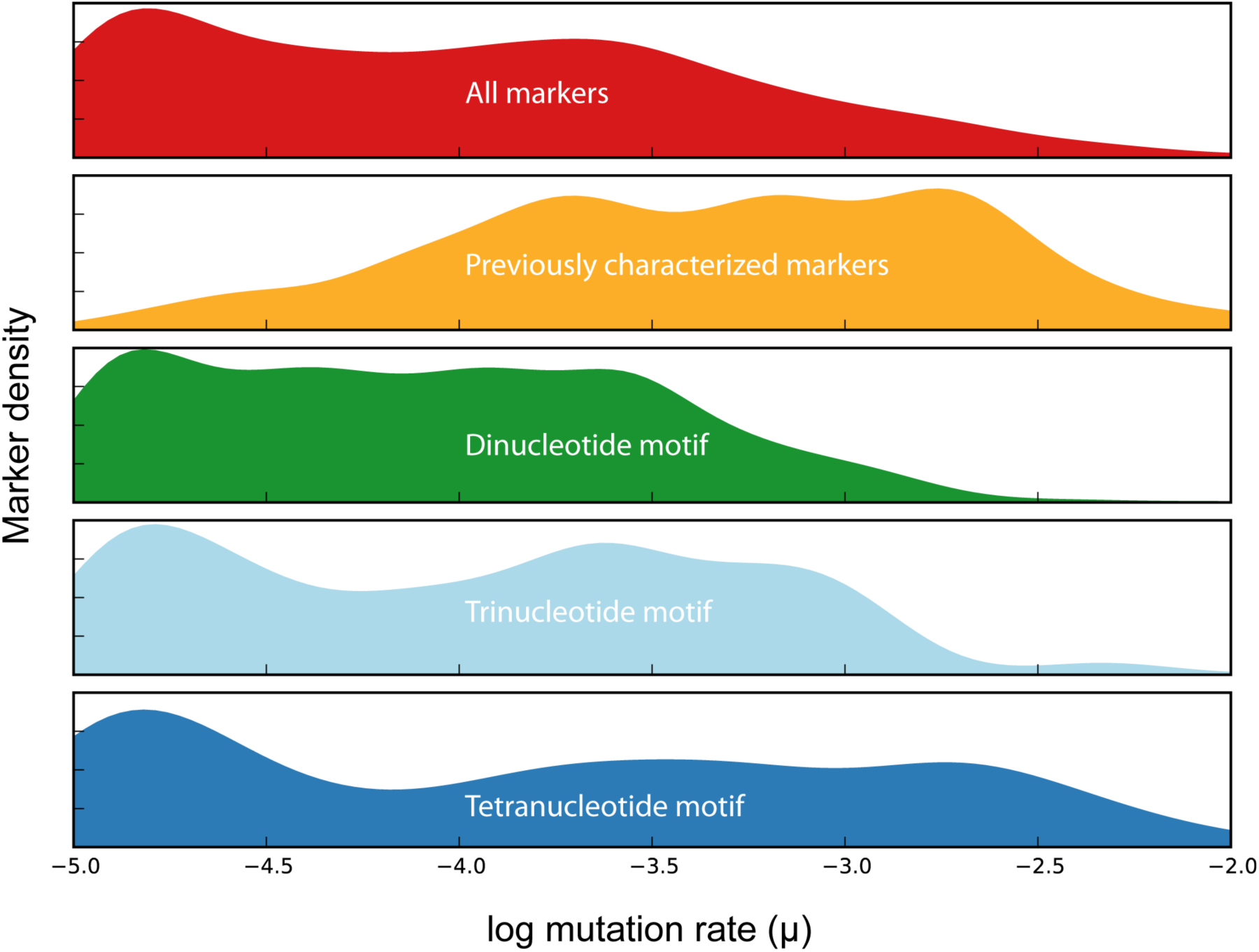
Distribution of Y-STR mutation rates. In red, we show the distribution of mutation rates across all STRs in this study. The set of loci with previously characterized mutation rates (orange) is substantially enriched for more mutable loci. When stratified by motif length, loci with tetranucleotide motifs (dark blue) are the most mutable, followed by loci with trinucleotide (light blue) and dinucleotide (green) motifs.

Leveraging our Y-STR mutation rate catalog, we searched for loci with relatively high mutation rates. These loci help to distinguish Y-chromosomes of highly related individuals and can help to precisely date patrilineal relatedness among individuals, which is important for forensics and genetic genealogy. Most of the markers with the greatest estimated mutation rates have been characterized in prior studies (**Table 1**), but we identified six loci whose mutation rates were estimated to be greater than ~2×10^−3^ mpg and are yet to be reported (**Tables 2-3**). Two of these markers, DYS548 and DYS467, have been used in previous genealogical panels but to the best of our knowledge, their mutation rates were never reported. In addition, we identified more than 65 loci with dinucleotide motifs and mutation rates greater than ~3.33×10^−4^ mpg (**Table 3, Table S1**).

**Table 1.** The most mutable Y-STRs with previously characterized mutation rates.

| Chrom | Hg19 start | Hg19 end | Motif | Mutation rate (mpg) | Homogeneous tract length (bp) | Name |
| --- | --- | --- | --- | --- | --- | --- |
| Y | 7,053,359 | 7,053,426 | AAAG | $1.37 \times 10^{-2}$ | 68 | DYS576 |
| Y | 7,867,880 | 7,867,943 | AAAG | $9.20 \times 10^{-3}$ | 64 | DYS458 |
| Y | 6,861,231 | 6,861,298 | AAAG | $7.80 \times 10^{-3}$ | 72 | DYS570 |
| Y | 14,515,312 | 14,515,363 | AGAT | $5.08 \times 10^{-3}$ | 48 | DYS439 |
| Y | 8,426,378 | 8,426,443 | AAG | $4.67 \times 10^{-3}$ | 69 | DYS481 |
| Y | 21,520,224 | 21,520,275 | AGAT | $4.50 \times 10^{-3}$ | 48 | DYS549 |
| Y | 18,718,889 | 18,718,940 | AGAT | $4.20 \times 10^{-3}$ | 52 | Y-GATA-A10 |
| Y | 4,270,960 | 4,271,019 | AGAT | $3.77 \times 10^{-3}$ | 60 | DYS456 |
| Y | 19,372,273 | 19,372,328 | AGAT | $2.88 \times 10^{-3}$ | 48 | DYS543 |
| Y | 14,761,101 | 14,761,160 | AGAT | $2.65 \times 10^{-3}$ | 46 | DYS442 |

**Table 2.** The most mutable Y-STRs with tetranucleotide motifs and previously uncharacterized mutation rates.

| Chrom | Hg19 start | Hg19 end | Motif | Mutation rate (mpg) | Homogeneous tract length (bp) | Name |
| --- | --- | --- | --- | --- | --- | --- |
| Y | 14,612,456 | 14,612,520 | AGAT | $5.07 \times 10^{-3}$ | 59 | DYS467 |
| Y | 5,409,729 | 5,409,801 | AAAG | $5.06 \times 10^{-3}$ | 61 | N/A |
| Y | 19,500,594 | 19,500,656 | AAAG | $4.89 \times 10^{-3}$ | 63 | N/A |
| Y | 14,200,743 | 14,200,802 | AGAT | $4.54 \times 10^{-3}$ | 56 | N/A |
| Y | 21,665,702 | 21,665,764 | AAAT | $3.66 \times 10^{-3}$ | 50 | DYS548 |

**Table 3.** The most mutable Y-STRs with dinucleotide motifs and previously uncharacterized mutation rates.

| Chrom | Hg19 start | Hg19 end | Motif | Mutation rate (mpg) | Homogeneous tract length (bp) | Name |
| --- | --- | --- | --- | --- | --- | --- |
| Y | 2,807,025 | 2,807,064 | AT | $3.62 \times 10^{-3}$ | 44 | N/A |
| Y | 2,708,412 | 2,708,457 | AG | $1.75 \times 10^{-3}$ | 46 | N/A |
| Y | 3,832,234 | 3,832,278 | AC | $1.66 \times 10^{-3}$ | 45 | N/A |
| Y | 6,398,638 | 6,398,684 | AC | $1.62 \times 10^{-3}$ | 49 | N/A |
| Y | 17,109,092 | 17,109,141 | AC | $1.57 \times 10^{-3}$ | 48 | N/A |

We observed wide variability in the mutation rates and patterns between motif length classes. STRs with tetranucleotide motifs had the greatest median mutation rate (*μ*=1.76×10^−4^ mpg), followed by those with trinucleotide (*μ*=1.22×10^−4^ mpg), pentanucleotide (*μ*=1.19×10^−4^ mpg), dinucleotide (*μ*=7.7×10^−5^ mpg), and hexanucleotide motifs (*μ*=3.28×10^−5^ mpg) (**Figure 4**). However, within each motif class, mutation rates varied by two or more orders of magnitude, indicating that other factors contribute to STR variability and highlighting that aggregate mutation rate statistics depend on the set of loci under consideration. We also found marked differences in the mutation patterns between motif classes. Loci with dinucleotide motifs and mutation rates greater than 10^−4^ mpg had a median step size parameter of *ρ*_*M*_ = 0.8, implying that many of the de novo mutations are expected to be greater than one repeat unit. On the other hand, the median step size parameter for longer motif classes within this mutation rate range was closer to one, implying that nearly all de novo events involve single step mutations.

Next, we harnessed the large number of Y-STR mutation rate estimates to identify the sequence determinants of mutation rates. For STRs without repeat structure interruptions, the length of the major allele explains a substantial fraction of the variance in log mutation rates for loci with di-, tri-, and tetranucleotide motifs (*R*^*2*^ = 0.83, *R*^*2*^ = 0.67, and *R*^*2*^ = 0.82, respectively; pentanucleotide motifs were not assessed due to a small number of data points). However, when analyzing all STRs, including those with interruptions, the length of the major allele is a poor predictor that explains only a modest amount of the variance (*R*^*2*^ = 0.16, *R*^*2*^ = 0.25, and *R*^*2*^ = 0.42) **(Figure 5, left panels**). To construct an improved model, we analyzed the relationship between the log mutation rate and the length of the longest uninterrupted repeat tract, regardless of the number of interruptions (**Figure 5, right panels**). This model explained more than 75% of the variance in mutability for each of the three motif length classes. To assess the impact of the repeat motif on the mutation rate, we stratified loci with dinucleotide motifs by repeat sequence (AC, AG, or AT) and once again regressed the log mutation rate on the length of either the major allele or longest uninterrupted tract (**Figure S9**). Major allele length was again a relatively poor predictor of the log mutation rate, but uninterrupted tract length explained more than 80% of the variance for each motif. Although these motif-specific models improved the *R*^*2*^, the increase was quite limited, suggesting that conditioned on the uninterrupted tract length, the repeat motif itself plays a minor role in the mutation rate. Taken together, our results show that a simple model of motif size and longest uninterrupted tract length largely explains STR mutation rates.

**Figure 5:**
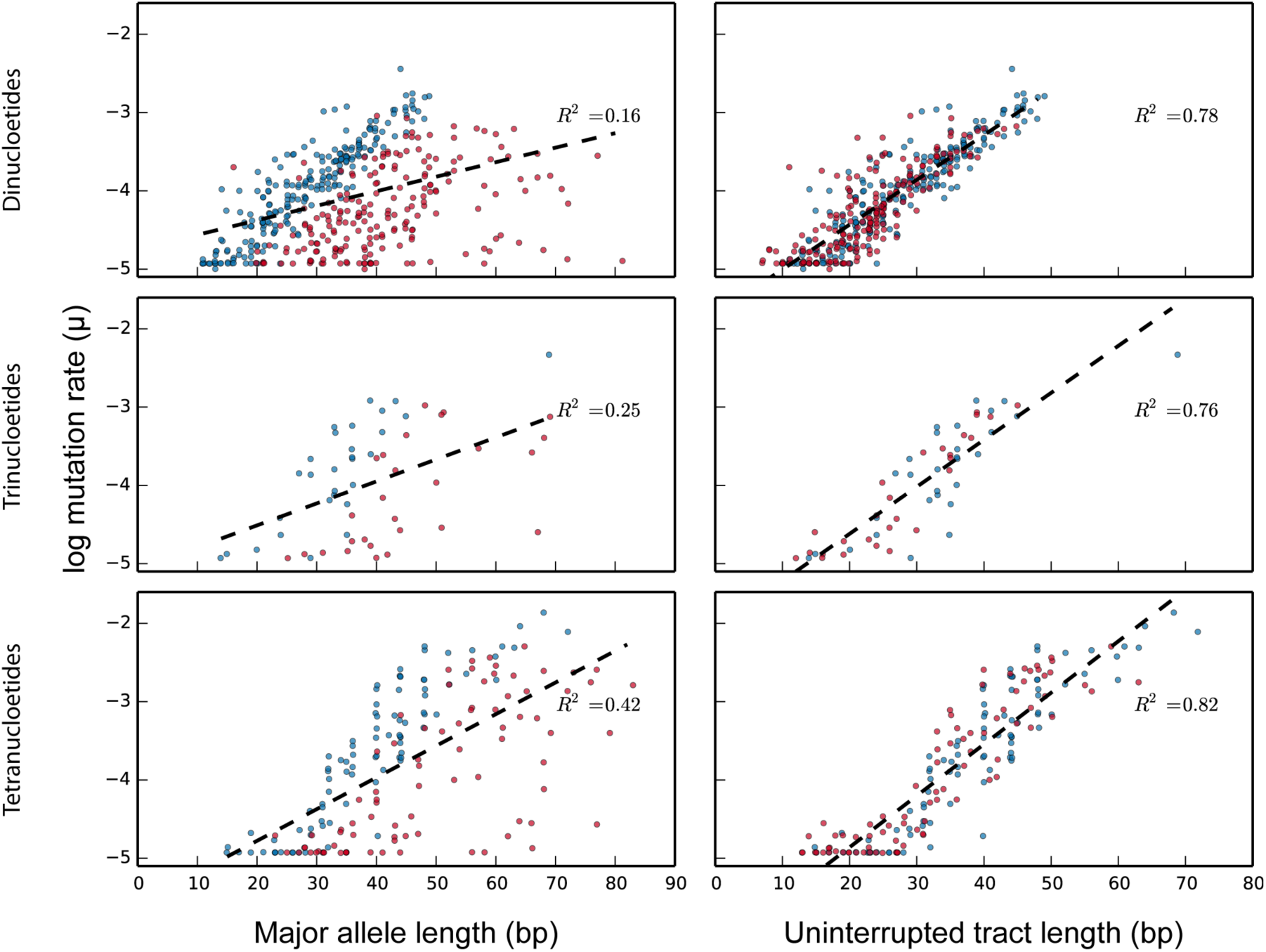
Sequence determinants of Y-STR mutability. Each panel presents the estimated log mutation rates (*y*-axis) of STRs versus either the major allele length (left panels, *x*-axis) or the longest uninterrupted tract length (right panels, *x*-axis) for various repeat motif sizes (rows). The black lines represent the mutation rate predicted by a simple linear model. For a given allele length (left panels), *Y*-STRs with no interruptions to the repeat structure (blue) are generally more mutable than those with one or more interruptions (red). Whereas major allele length alone is poorly correlated with mutation rate (left panels), the longest uninterrupted tract length (right panels) is strongly correlated regardless of the number of interruptions.

### Predicting Genome-Wide STR Mutation Rates

We estimated the number of de novo mutations across the entire genome using the determinants found above. For each repeat motif length, we trained a non-linear mutation rate predictor using the uninterrupted tract lengths and mutation rates of the polymorphic Y-STRs. To account for the fixed STRs in our dataset and to better fit the model at shorter tract lengths, we assigned each fixed locus a mutation rate of 10^−5^ mpg, the lower mutation rate boundary used by MUTEA (**Figure S10**), and we jointly trained the predictors across all STRs. To validate these predictors, we used them to estimate the mutation rates of paternally transmitted autosomal CODIS markers, which the National Institute of Standards and Technology (NIST) has previously estimated using conventional means. Our predictors explained about 75% of the variance in the log mutation rates for these markers. In addition, the median mutation rate reported by NIST (*μ*=1.3×10^−3^ mpg) closely matched the result reported by our predictors (*μ*=1.0×10^−3^ mpg), suggesting that they generate reliable predictions.

Next, we ran our predictors on each STR in the human genome with 2-4 bp motifs, resulting in mutation rate estimates for each of the ~590,000 loci (**Table S2**). Since our model was trained using Y-STR mutation rates, these estimates refer only to the paternally inherited half of the genome. We discarded estimated rates below 1.25×10^−5^ mpg, as these are too close to the MUTEA lower boundary and may therefore be upwardly biased. After filtering, our model predicts that there are ~70,000 STRs with mutation rates greater than 10^−4^ mpg, ~44,000 loci with mutation rates greater than 1 in 3000 mpg and that an STR should mutate at an average rate of 4.4×10^−4^ mpg. Stratifying our results by motif length, we predict 29, 3, and 33 de novo STR mutations for loci with di-, tri- and tetranucleotide motifs on the paternally inherited set of chromosomes.

Overall, we predict that 76-85 de novo STR mutations occur each generation for the full set of chromosomes. To account for the maternal chromosomes, we extrapolated our paternal results using prior estimates of the male to female STR mutation rate ratio (3.3:1 to 5.5:1^19; 50^). We posit that our estimates for STR de novo mutational load are likely to be conservative. First, we omitted loci with 5-6 bp motifs for which we did not have sufficient data to build a mutation rate model. Second, for autosomal STRs whose uninterrupted tract lengths exceeded the maximal length observed in our study, we estimated their mutation rates using the maximal Y-STR length. Given the strong positive correlation between tract length and mutation rate observed in our study, these loci are probably far more mutable. Despite our conservative approach, the estimated number of genome-wide de novo STR mutations rivals that of any known class of genetic variation, including SNPs (~70 events per generation), indels (1-3 events), and SV and interspersed repeats (<1 event per generation).^6; 7; 9; 51^ As such, our results highlight the putative contribution of STRs to de novo genetic variation.

### Y-STRs in Forensics and Genetic Genealogy

We assessed the applicability of our Y-STR results to the genetic genealogy and forensic DNA communities. First, we considered whether it would be possible to distinguish between closely patrilineally related individuals from high-throughput sequencing data. Based on the entire Y-STR set reported by our study, we expect roughly one de novo mutation to occur every four generations. In addition, from WGS data, one also expects to identify approximately one de novo SNP every 2.85 generations^35^, resulting in a 60% theoretical probability of differentiating between a father and son’s Y-chromosome haplotype using high-throughput sequencing. Previous studies have suggested that capillary genotyping of 13 rapidly mutating Y-STRs can discriminate between father-son pairs in 20-27% of the cases.^21; 52^ However, these particular markers are largely inaccessible to whole-genome sequencing data due to their long length and highly repetitive flanking regions that preclude unique mapping. With increased interest in high-throughput sequencing among genetic genealogy services (e.g. FullGenomes and Big Y by FamilyTreeDNA) and the forensics community, our results suggest that WGS can achieve better patrilineal discrimination compared to common panel-based methods. Of course, the main caveat is that WGS technology is at least an order of magnitude more expensive than a panel-based approach. However, if the current trajectory of sequencing cost decline continues, shotgun sequencing to discriminate between closely patrilineally related individuals might soon become economically viable.

We also assessed the accuracy of imputing Y-STR profiles from Y-SNP data. This capability may be useful in forensic cases involving a highly degraded male sample for which complete Y-STR profiles would be difficult to obtain. In such cases, since there are many more SNPs than STRs on the Y-chromosome, it might be possible to salvage some of those markers with a high-throughput method and impute Y-STRs profiles for compatibility with common forensic or genealogical databases.

For imputation, we created a framework called MUTEA-IMPUTE. Briefly, after building a SNP phylogeny relating all samples and learning a mutation model as outlined in **Figure 1**, MUTEA-IMPUTE passes two sets of messages along the phylogeny to compute the exact marginal posteriors for each node, resulting in imputation probabilities for samples without observed Y-STR genotypes (**Appendix D**). We assessed the accuracy of our algorithm by imputing the 1000 Genomes individuals for the PowerPlex Y23 panel, a set of markers regularly used in forensic cases involving sex crimes. Over 100 iterations, we randomly constructed reference panels of 500 samples and used MUTEA-IMPUTE to calculate the maximum a posteriori genotypes for a distinct set of 70 samples.

Despite the small size of the reference panel, we were able to correctly impute an average of 66% of the genotypes without any quality filtration (**Table S3**). Importantly, the resulting imputed probabilities roughly matched the average accuracy, indicating that the posteriors computed using this technique are well calibrated (**Figure S11**). Discarding imputed genotypes with posteriors below 70% resulted in an overall accuracy of 88% and retained about 40% of the calls. On a marker-by-marker basis, accuracy was generally inversely proportional to the estimated mutation rates, with the most slowly mutating markers having accuracies on the order of 95%. This trend stems from the fact that as the mutation rate increases, shorter branch lengths are required to obtain an estimate with similar confidence. We envision that a larger panel will substantially increase the ability to correctly impute Y-STRs and might facilitate work with highly degraded samples, a common issue in forensics casework.

## Discussion

Advances in sequencing technology have fundamentally altered Y-STR analyses. The initial scarcity of SNP genotypes led to the development of methods to infer coalescent models from Y-STR genotypes alone. Methods designed to also learn STR mutational dynamics either marginalized over these coalescent models^53^ or aimed to simultaneously infer the coalescent and mutational models.^54; 55^ With the advent of population-scale WGS datasets, many of these STR-centric approaches have instead used SNPs, resulting in substantially more detailed phylogenies. For the Y-chromosome, these detailed phylogenies now provide the evolutionary context required to interpret Y-STR mutations, obviating the need for computationally expensive tree enumeration or marginalization approaches. However, the errors prevalent in WGS-based Y-STR genotypes require methods capable of accounting for genotype uncertainty, precluding the application of many traditional microsatellite distance measures designed for capillary data.^44; 45^

In this study, we developed MUTEA, a method that leverages population-scale sequencing data to estimate Y-STR mutation rates. One inherent advantage of our approach is its ability to model and learn many of the salient features of microsatellite mutations. By incorporating a geometric step-size distribution, we allow both single-step mutations that predominate at tetranucleotide loci^19; 56^ as well as multistep mutations that frequently occur at dinucleotide loci.^19; 57^ In addition, the model’s length constraint parameter captures the intra-locus phenomenon of shorter STR alleles preferentially expanding and longer alleles preferentially contracting.^57; 58^ As these parameters are learned from observed STR genotypes, our method avoids many biases that stem from imposing single-step mutations or assuming parameters a priori.

In addition to its mutational model flexibility, our approach has both high throughput and a high dynamic range. With whole-genome sequencing data, we were able to assess every Y-STR that is accessible to Illumina sequencing, dramatically increasing the catalog of polymorphic loci with estimated mutation rates. In addition, by leveraging deep Y-chromosome phylogenies, we were able to obtain mutation rate estimates for very slowly mutating loci. Our estimates were highly replicable and consistent, as demonstrated by the strong concordance between the estimates from the two whole-genome sequencing datasets.

Our approach has several inherent limitations. Because Illumina datasets are currently comprised of 75-100 base pair reads, we were unable to genotype and characterize the mutation rates of both long Y-STRs and Y-STRs that reside in heterochromatic regions. Due to the strong relationship between tract length and mutation rate, we anticipate that more rapidly mutating loci reside on the Y-chromosome. In addition, we were unable to characterize the mutation rates of homopolymers due to a rapid degradation of base quality scores with increasing allele length. As a result, future studies may benefit from reapplying our analyses as sequencing technologies, particularly those enabling longer reads, continue to mature. Another limitation is that our mutation model does not capture the full complexity of STR mutational dynamics, as it ignores intra-locus mutation rate variation.^59^ Incorporating these and other mutational characteristics may be of interest to future studies.

One longstanding question regarding Y-STR mutation rates has been the apparent discrepancy between evolutionary and pedigree-based mutation rates. Several studies have suggested that evolutionary rates are 3-4 times lower, resulting in substantial inconsistencies in Y-STR-based lineage dating and large discrepancies from Y-SNP-based TMRCA estimates.^20; 46; 60^ Because our study harnessed evolutionary data, we sought to avoid any potential issues by scaling each phylogeny such that our estimates best matched those from pedigree-based studies. Nonetheless, our investigations into an alternative scaling based on a SNP molecular clock resulted in similar scaling factors that only differed by ~25%. Coupled with the strong concordance we observed with pedigree-based estimates, our study provides little evidence for a substantial difference between mutation rates estimated from these two types of data. Future work may benefit from assessing whether these previously reported discrepancies were due to the simplified Y-STR mutation models used in the approaches to obtain evolutionary-based Y-STR mutation rates.

Our large corpus of mutation rate estimates has enabled us to dissect the sequence factors governing STR mutability. We determined that the longest uninterrupted tract length is a strong predictor of the log mutation rate. This observation matches the exponential relationship between mutation rate and tract length previously reported in several pedigree-based studies.^21; 50; 56; 58^ We also found that the total length of the major allele was a poor predictor. Coupled with the fact that Y-STRs without interruptions were much more mutable than interrupted ones with the same major allele length, our study provides strong evidence that interruptions to the repeat structure decrease mutation rates. This finding supports what has long been posited in STR evolutionary models^61; 62^ and has been shown in a handful of small-scale experimental studies of STR mutability.^63; 64^ However, it contradicts the recent findings of Ballantyne et al. in which no effect was observed.^21^

Another open question is why STRs with dinucleotide motifs have lower mutation rates, given their higher levels of polymorphisms in the population. A previous large-scale panel-based study reported that loci with dinucleotide motifs have lower mutation rates than loci with tetranucleotide motifs.^19^ Our survey confirmed this observation without ascertainment of STRs directly based on their polymorphism rates. However, genome-wide analyses of STRs have shown that dinucleotides have more diverse allelic spectra than tetranucleotides.^23; 26^ These results are unlikely to be due to genotyping errors as a study of an individual sequenced to a depth of 120× also showed that dinucleotide repeats are more polymorphic than other types of STRs.^23^ One potential explanation is that STRs with dinucleotide motifs have larger step sizes but lower mutation rates. However, we cannot exclude other explanations such as a difference in length constraint.

Our large compendium of mutation rate estimates has also enabled predictions about genome-wide STR variation. Prior studies have estimated a rate of approximately 75 de novo mutations per generation^4; 8^ but have largely ignored STRs, despite their elevated mutation rates. Based on our projections for paternally inherited chromosomes, the number of de novo STR mutations is likely to rival the combined contribution of all other types of genetic variants. As several lines of evidence have highlighted the involvement of STR variations in complex traits^11–13; 65^, it will be important to assess the biological impact of these de novo STR variations on human phenotypes.

## Appendix A. Simulating Exact STR Genotypes

Values of *μ*, *β*, and *ρ*_*M*_ ranging from 10^−5^ to 10^−2^, 0 to 0.5, and 0.75 to 1.0, respectively, were used to simulate genotypes under a wide range of mutation models. Using either the 1000 Genomes phylogeny or the SGDP phylogeny, each simulation was performed as follows:

1. Randomly assign the root node an STR allele between −4 and 4 and mark it as active
2. Remove an active node and mark it as inactive. For each of this node’s children:

i. Calculate the child’s allele probabilities using the branch length, the true mutation model and the parent node’s genotype
ii. Randomly select an STR allele based on these probabilities
iii. Mark the descendant node as active
3. While active nodes remain, go to step 2
4. Report the exact STR alleles for a random subset of the samples (leaf nodes) based on the required sample size

## Appendix B. Appendix B. Simulating STR Sizes in Reads with PCR Stutter

We first used the procedure above to simulate STR genotypes down the phylogeny. We then used the true genotype for a particular sample *g*_*i*_ and a given stutter model to simulate the STR sizes observed in each read as follows:

1. Sample the number of observed reads *n*_*reads*,*i*_; for each sample with genotype *g*_*i*_ from the read count distribution
2. For each read from 1 through *n*_*reads,i*_, sample a number *c* ~ U (0,1)

i. If *c* < *d*, randomly sample an artifact size *a*_*j*_ from a geometric distribution with parameter *ρ*_*s*_. Report the read’s STR size as *g*_*i*_ – *a*_*j*_
ii. If *d* ≤ *c* < 1 – *u*, report the read’s STR size as *g*_*i*_
iii. Otherwise, randomly sample an artifact size *a*_*j*_ from a geometric distribution with parameter *ρ*_*s*_. Report the read’s STR size as *g*_*i*_ + *a*_*j*_

To assess whether estimates would be accurate for even the most sparsely sequenced loci, we used read count distributions obtained from both Y-STR call sets corresponding to loci in the 10^th^ coverage percentile. For **Figure 2**, we used a stutter model with *d* = 0.15, *u* = 0.01 and *ρ*_*s*_ = 0.8, and we used 1, 2 and 3 reads for 65%, 25% and 10% of samples, respectively.

## Appendix C. Confidence Interval Estimation

We used a delete-*d* jackknife approach to estimate mutation rate confidence intervals.^66^ For each Y-STR, we sampled without replacement half of the STR genotypes a total of 100 times and estimated the log mutation rate using each of these subsets. Given these subsample estimates and the log estimate obtained using all samples, the standard error (SE) and confidence interval (CI) for the log mutation rate were calculated according to:

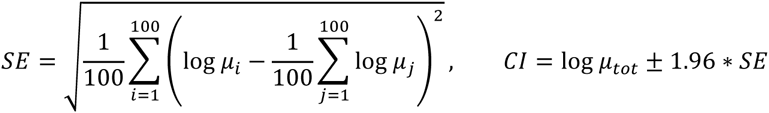

where *μ*_tot_ is the estimate based on the full dataset.

## Appendix D: Y-STR Imputation

We extended MUTEA to impute missing STR genotypes. Using the approach outlined in **Figure 1**, we first construct a phylogeny relating all samples and learn a mutation model. We then use this learned mutation model to pass two sets of messages along the tree and compute exact posteriors for each node, resulting in imputation probabilities for samples with missing genotypes. For node *N*_*i*_ with parent *P*_*i*_, sibling *C*_1*i*_ and *C*_2*i*_ and children *S*_*i*_ and *C2Ì*, its conditional genotype probability given the observed data *D* is:

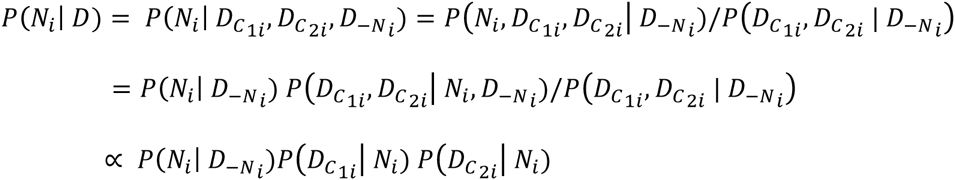

Here, *D*_*N*_*i*__ and *D*_–*N*_*i*__ denote the genotype likelihoods in and not in node *N*_*i*_’s subtree, respectively. We note that each of these terms is conditioned on the STR mutational model *M* and the Y-chromosome phylogeny *T*, but we omit these terms here and below for brevity.

The second and third terms in the node posterior expression are computed using a bottom-up traversal of the tree from the leaves to the root node. Each node in the tree combines information from its two children using the recurrence

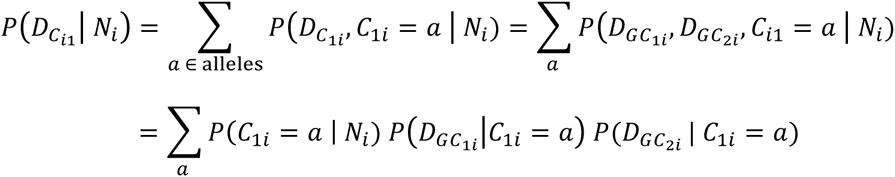

Here, *GC*_1*i*_ and *GC*_2*i*_ denote the two children of node *C*_1*i*_. This recurrence applies to all nodes except the leaves, where genotype posteriors or a uniform prior are used for samples with and without genotype information, respectively.

Similarly, the first term in the node posterior expression is computed using a top-down traversal of the tree from the root to the leaves. After assigning the root node a uniform prior probability, each node combines information from its parent and sibling:

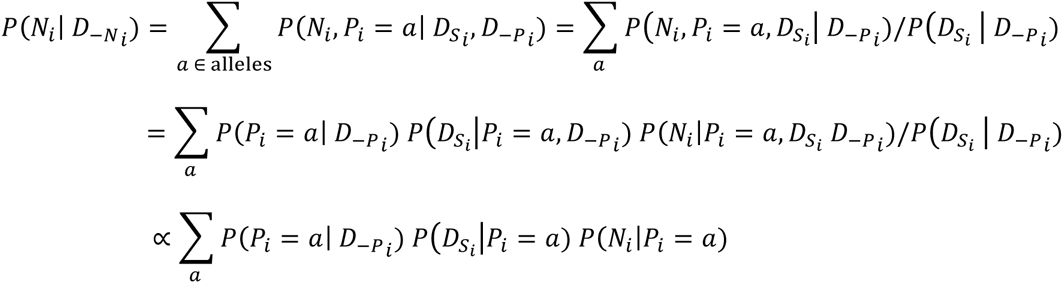

## Supplemental Data

Supplemental Data includes eleven figures and three tables.

## Acknowledgments

M.G. was supported by the National Defense Science and Engineering Graduate Fellowship. G.D.P. was supported by the National Science Foundation Graduate Research Fellowship under grant DGE-1147470. C.T.-S. was supported by The Wellcome Trust grant 098051. Y.E. holds a Career Award at the Scientific Interface from the Burroughs Wellcome Fund. This study was supported by NIJ grant 2014-DN-BX-K089 (Y.E. and T.W.). Y.E. is a SAB member of Identity Genomics, BigDataBio and Solve Inc. G.D.P is an employee of 23andMe. None of these entities played a role in the design, execution, interpretation, or presentation of this study.

## Web Resources

1000 Genomes Project BAM alignments,

ftp://ftp.1000genomes.ebi.ac.uk/vol1/ftp/data_collections/1000_genomes_project/data/

1000 Genomes Project capillary genotypes,

ftp://ftp.1000genomes.ebi.ac.uk/vol1/ftp/technical/working/20140107_chrY_str_haplotypes/YST

Rs_PowerPLexY23_1000Y_QA_20130107.txt

MUTEA, https://github.com/tfwillems/MUTEA

Dendroscope software, http://dendroscope.org/

HipSTR software, https://github.com/tfwillems/HipSTR

RAxML software, http://sco.h-its.org/exelixis/web/software/raxml/index.html

Simons Genome Diversity Project, https://www.simonsfoundation.org/life-sciences/simons-genome-diversity-project-dataset/

Simons Genome Diversity Project capillary genotypes, ftp://ftp.cephb.fr/hgdp_supp9/genotype-supp9.txt

Y-STR references, HipSTR call sets and Y-SNP phylogenies, https://github.com/tfwillems/ystr-mut-rates

**Figure S1:**
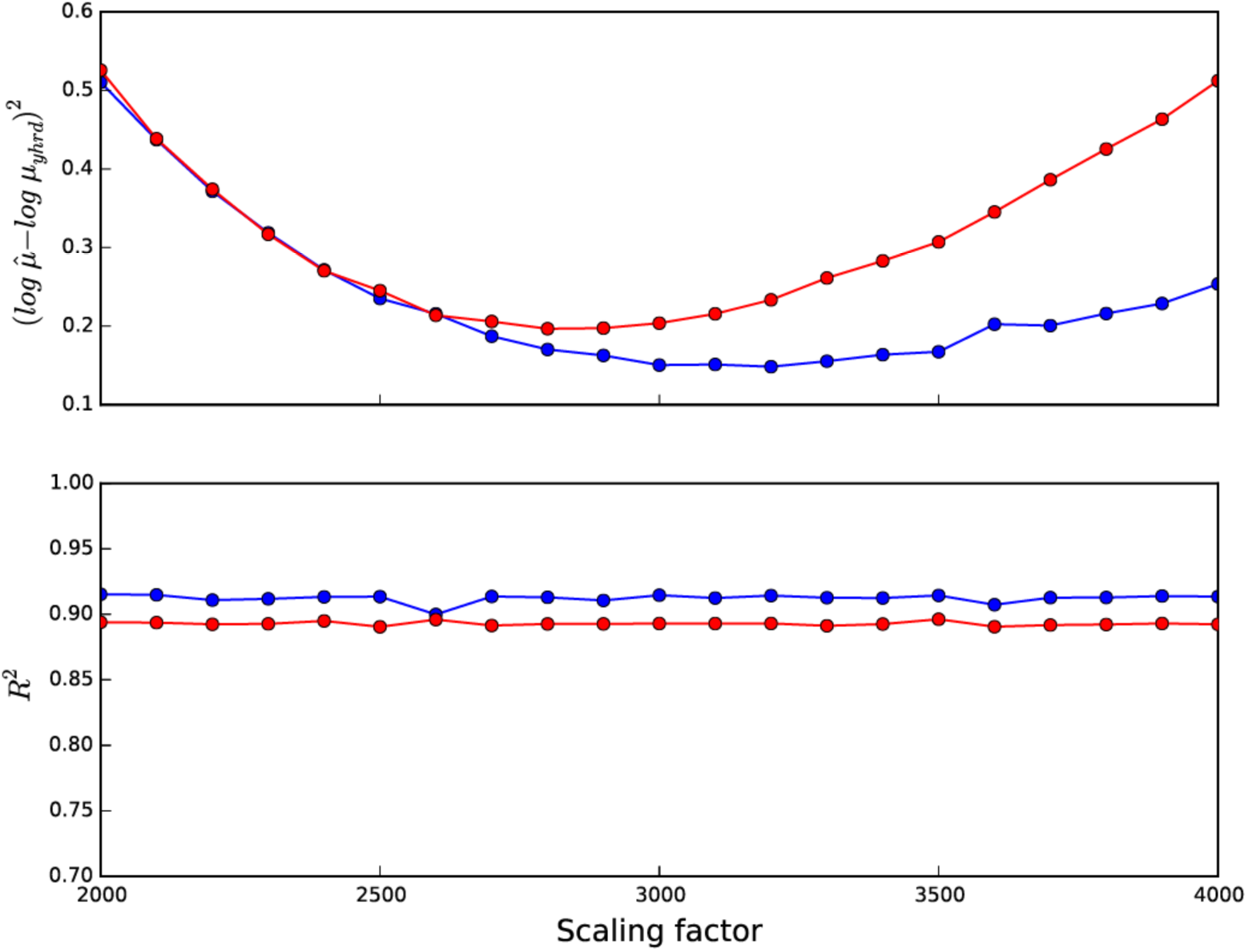
Scaling the Y-SNP phylogenies. Comparison of mutation rates for loci in the Y-Chromosome Haplotype Reference Database to estimates for the same loci obtained using data from the Simons Genome Project (blue) and the 1000 Genomes Project (red), over a range of scaling factors. Although the scaling factor had little effect on the *R*^2^, it substantially impacted the total squared error in the log estimates. For each data set, we selected the optimal scaling factor as the value that minimized this squared error.

**Figure S2:**
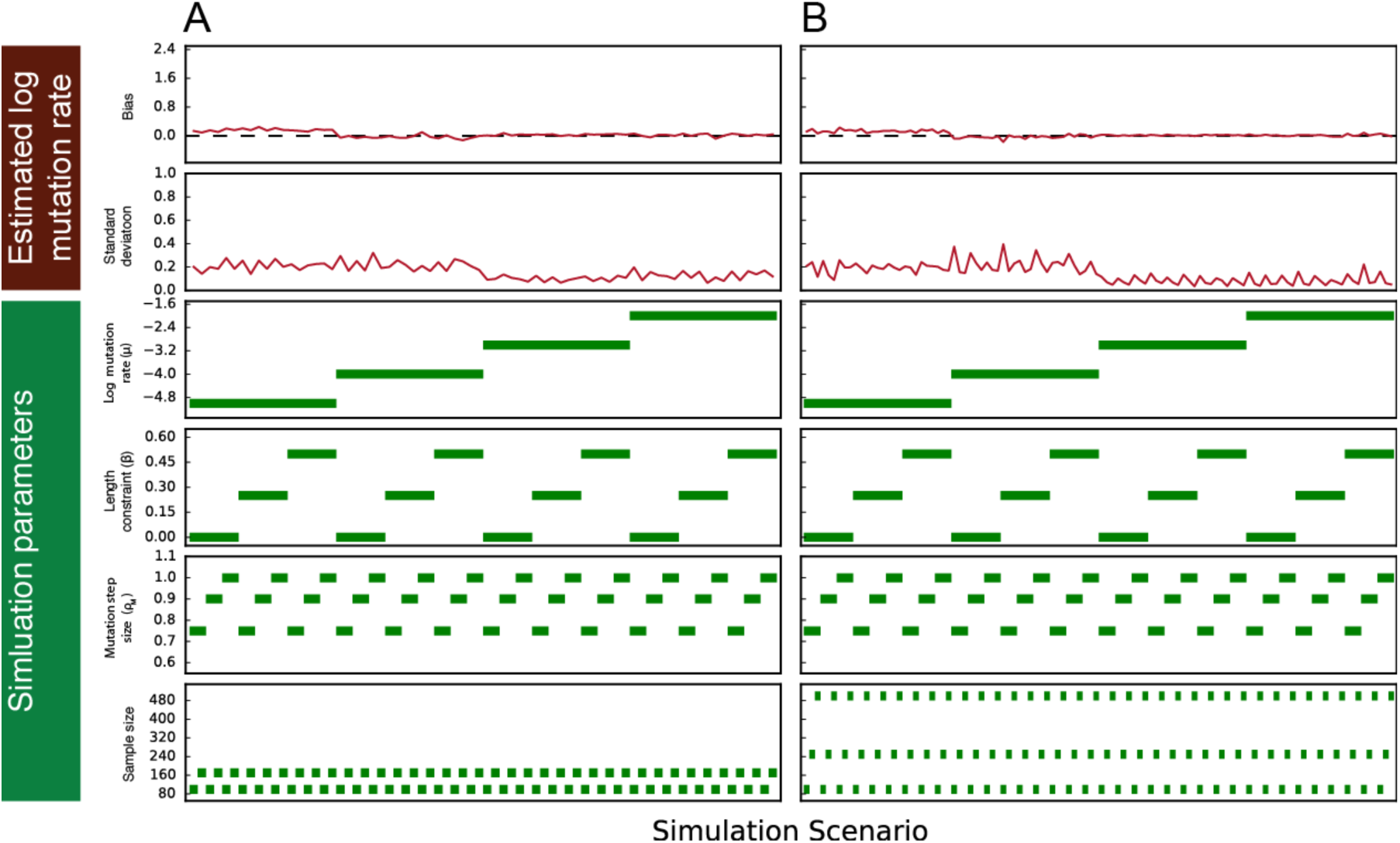
MUTEA obtains accurate mutation rate estimates from exact genotypes. STR genotypes were simulated for a variety of sample sizes and mutation models (four lower rows) for both the Simons Genomes phylogeny (**A**) and the 1000 Genomes phylogeny (**B**). After assigning each sample’s genotype a posterior probability of one and iterating 25 times for each simulation scenario, mutation rate estimates were unbiased (upper row) and have reasonably low standard deviations (second row).

**Figure S3:**
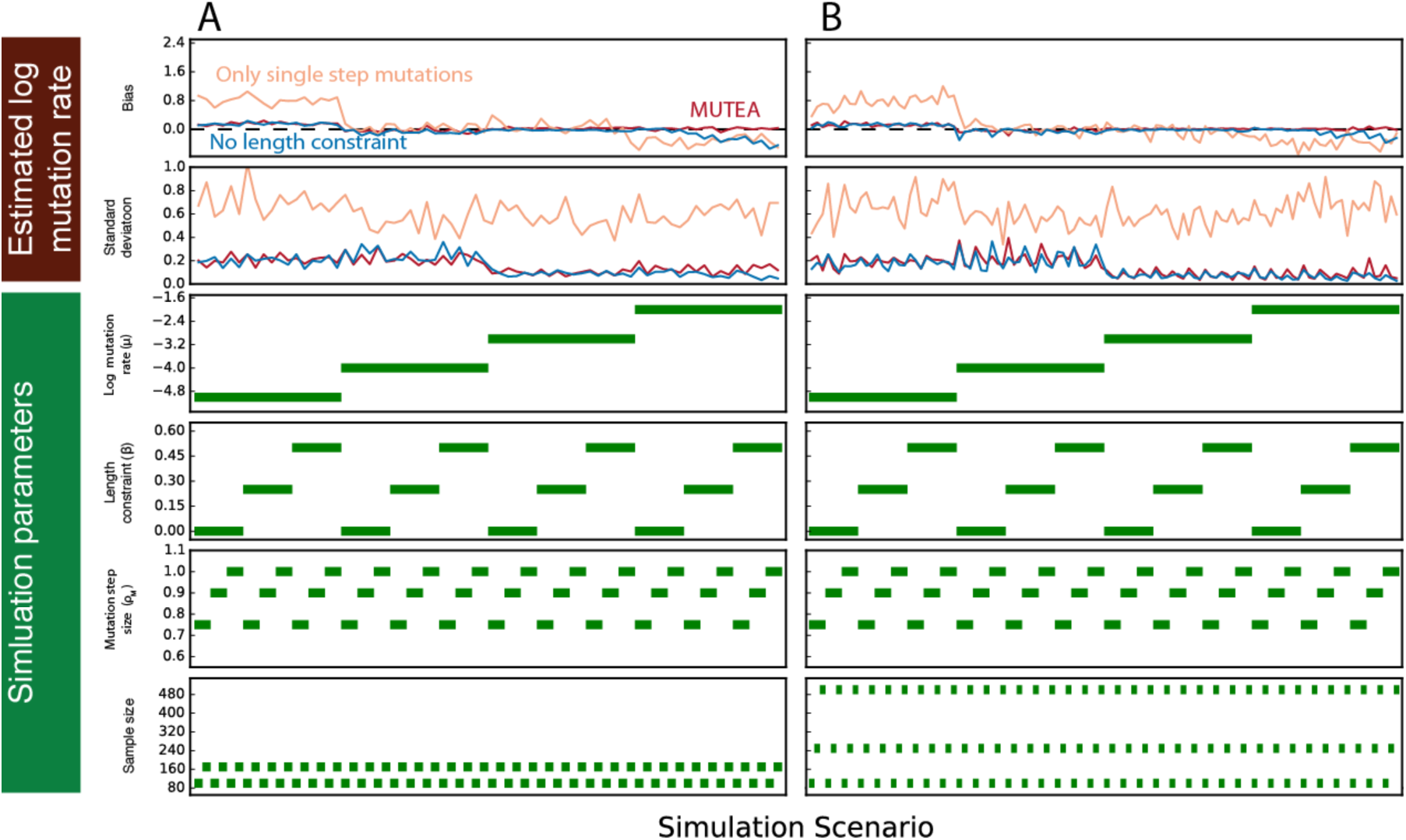
Simplifying mutation models results in biased mutation rate estimates. STR genotypes were simulated for a variety of sample sizes and mutation models (four lower rows) for both the Simons Genomes phylogeny (**A**) and the 1000 Genomes phylogeny (**B**). After assigning each sample’s genotype a posterior probability of one and iterating 25 times for each simulation scenario, mutation rate estimates were biased (upper row) when the estimated model is restricted to singe-step mutations (orange) or to no length constraint (blue), but not when the estimated model is unrestricted, as in MUTEA (red).

**Figure S4:**
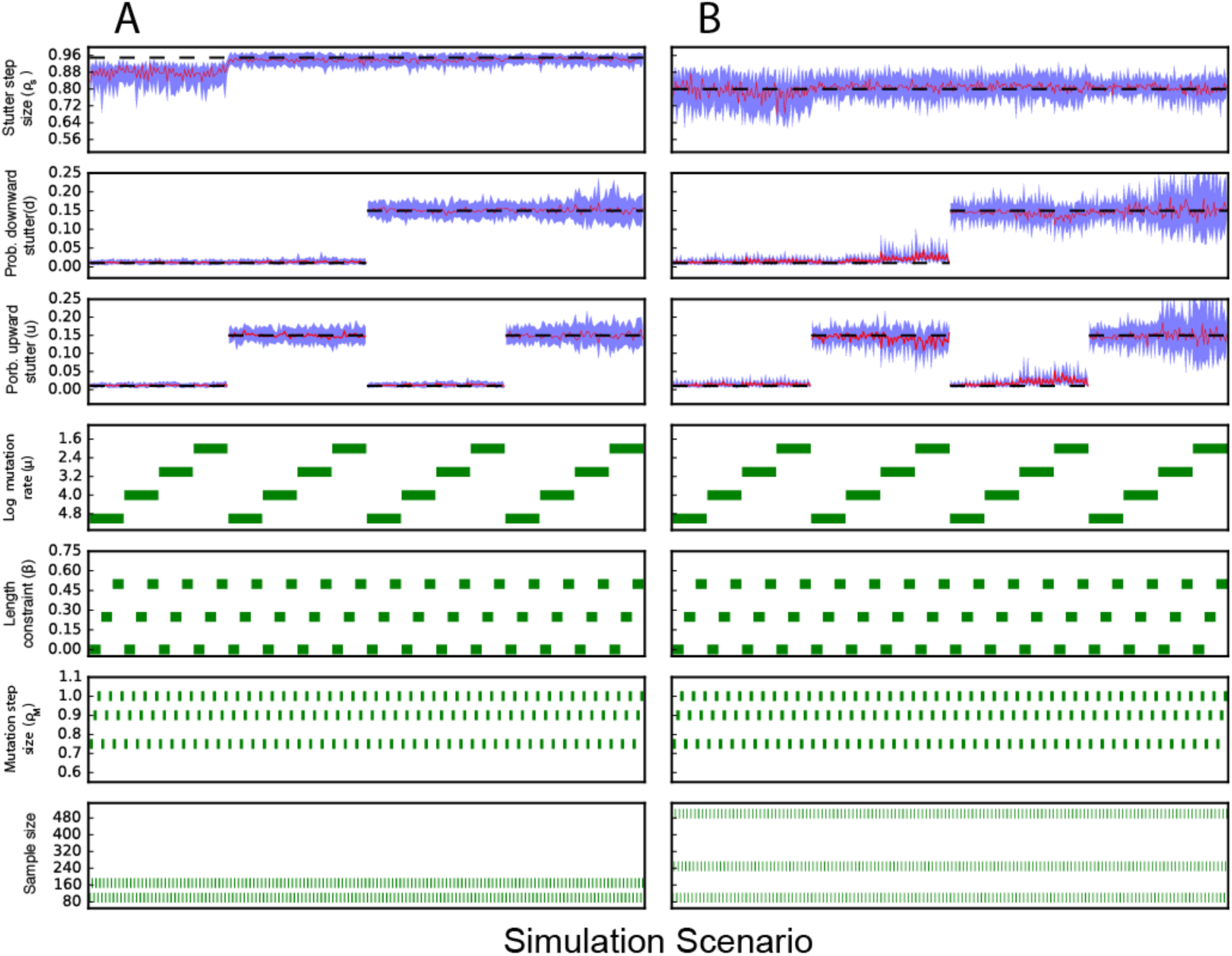
MUTEA accurately recovers the underlying stutter model. STR genotypes were simulated for a variety of sample sizes and mutation models (four lower rows). We then simulated observed reads for each set of genotypes using various PCR stutter models (dashed black lines in three upper rows) and input these simulated reads to MUTEA. Across 25 iterations of each scenario, the median inferred stutter parameters (red lines) were relatively unbiased. Blue lines indicate the lower and upper quartiles of the estimates for each scenario. **A**. 1, 2, 3, 4, 5 or 6 reads were generated for 19%, 27%, 21%, 15%, 8% and 10% of Simons Genome Diversity project samples, respectively. **B**. 1, 2 or 3 reads were generated for 65%, 25% and 10% of 1000 Genomes Project samples, respectively.

**Figure S5:**
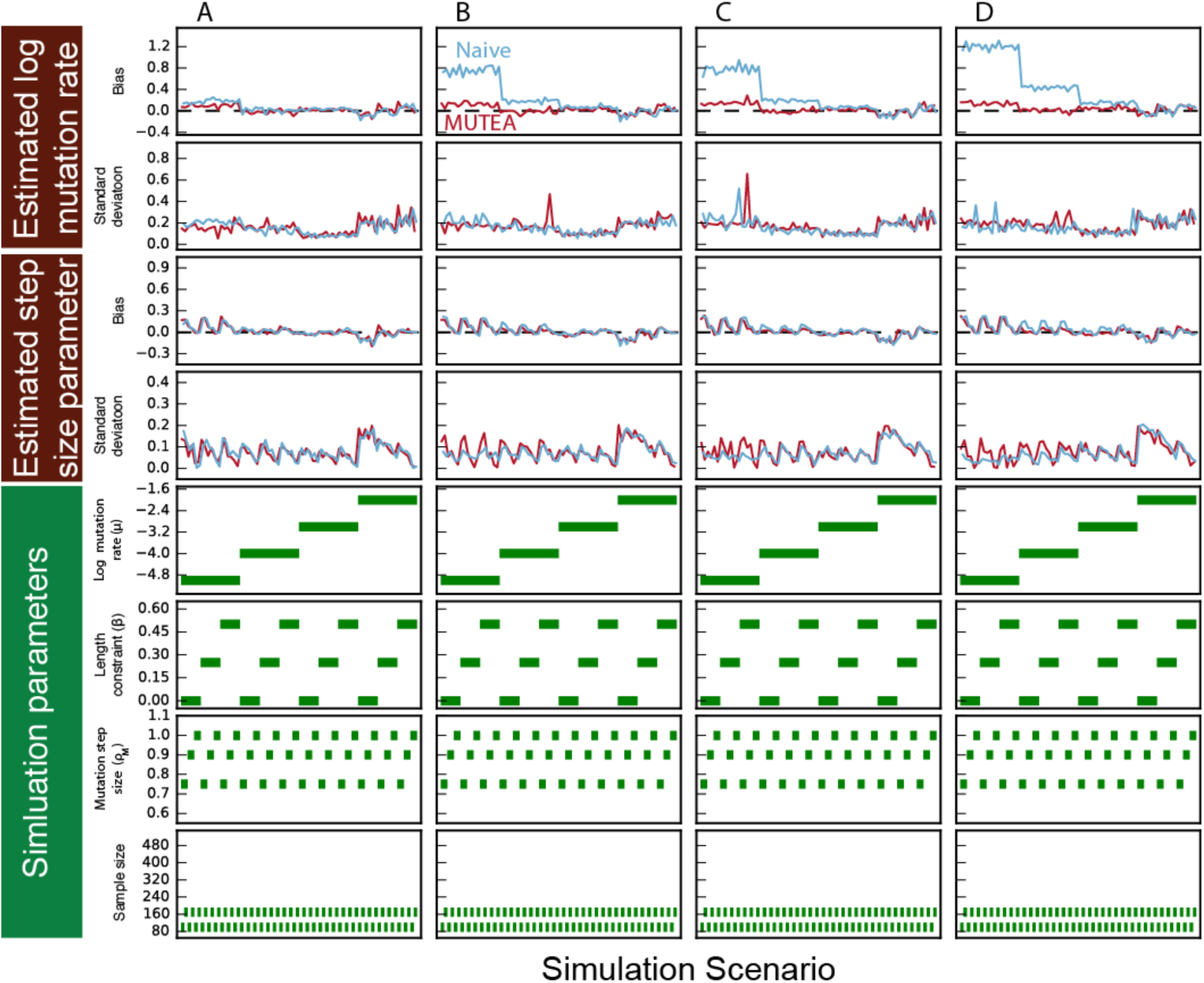
MUTEA infers unbiased mutation rates and step size parameters from stutter-affected reads using the Simons Genome Diversity Project phylogeny. STR genotypes were simulated for a variety of sample sizes and mutation models (four lower rows) using the Simons Genome phylogeny. Reads for each set of genotypes were then simulated using various PCR stutter models and input to MUTEA. Across 25 iterations for each scenario, MUTEA inferred unbiased estimates for the log mutation rate (red lines, upper row) and the step size parameter (third row). In contrast, a naïve method that computes genotype posteriors based on the fraction of supporting reads resulted in biased mutation rate estimates (blue lines). For each simulation, 1, 2, 3, 4, 5, or 6 reads were simulated for 19%, 27%, 21%, 15%, 8%, and 10% of samples using a stutter model with *ρ*_*s*_ = 0.95 and (**A**) *d* = 0.01 and *u* = 0.01, (**B**) *d* = 0.15 and *u* = 0.01, (**C**) *d* = 0.01 and *u* = 0.15, or (**D**) *d* = 0.15 and *u* = 0.15.

**Figure S6:**
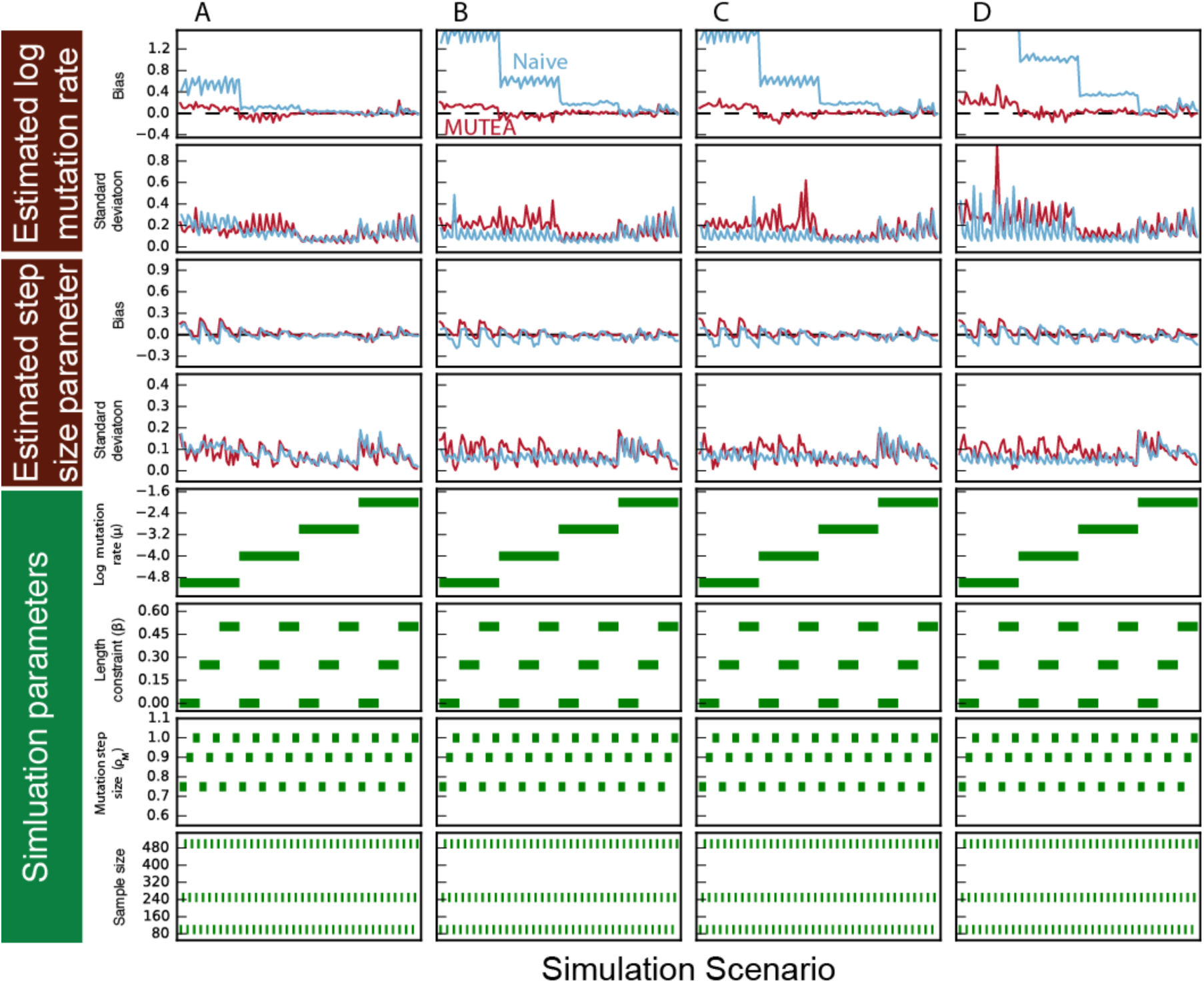
MUTEA infers unbiased mutation rates and step size parameters from stutter-affected reads using the 1000 Genomes Project phylogeny. STR genotypes were simulated for a variety of sample sizes and mutation models (four lower rows) using the 1000 Genome phylogeny. Reads for each set of genotypes were then simulated using various PCR stutter models and input to MUTEA. Across 25 iterations for each scenario, MUTEA inferred unbiased estimates for the log mutation rate (red lines, upper row) and the step size parameter (third row). In contrast, a naïve method that computes genotype posteriors based on the fraction of supporting reads resulted in biased mutation rate estimates (blue lines). For each simulation, 1, 2, or 3 reads were generated for 65%, 25%, and 10% of samples using a stutter model with *ρ*_*s*_ = 0.8 and (**A**) *d* = 0.01 and *u* = 0.01, (**B**) *d* = 0.15 and *u* = 0.01, (**C**) *d* = 0.01 and *u* = 0.15, or (**D**) *d* = 0.15 and *u* = 0.15.

**Figure S7:**
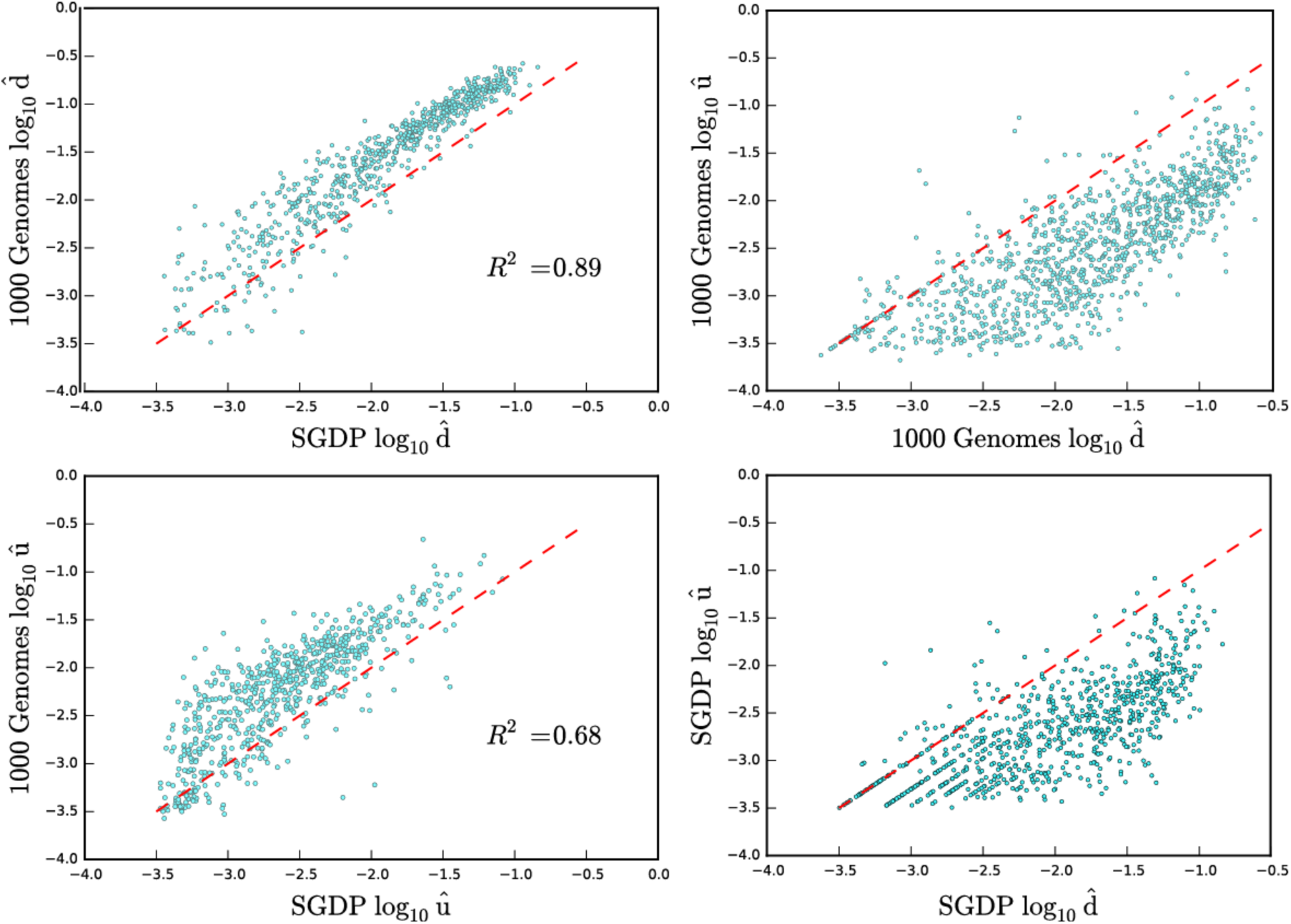
Relationships between stutter probabilities within and across datasets. For a given Y-STR locus, the probabilities of stutter increasing (*u*) or decreasing (*d*) the size of the STR in each read were highly correlated between input datasets (first column). However, the 1000 Genomes stutter rates largely fell above the diagonal (red line), indicating higher rates of stutter in this dataset. Within each dataset, nearly all loci had a higher rate of downward stutter than of upward stutter (second column).

**Figure S8:**
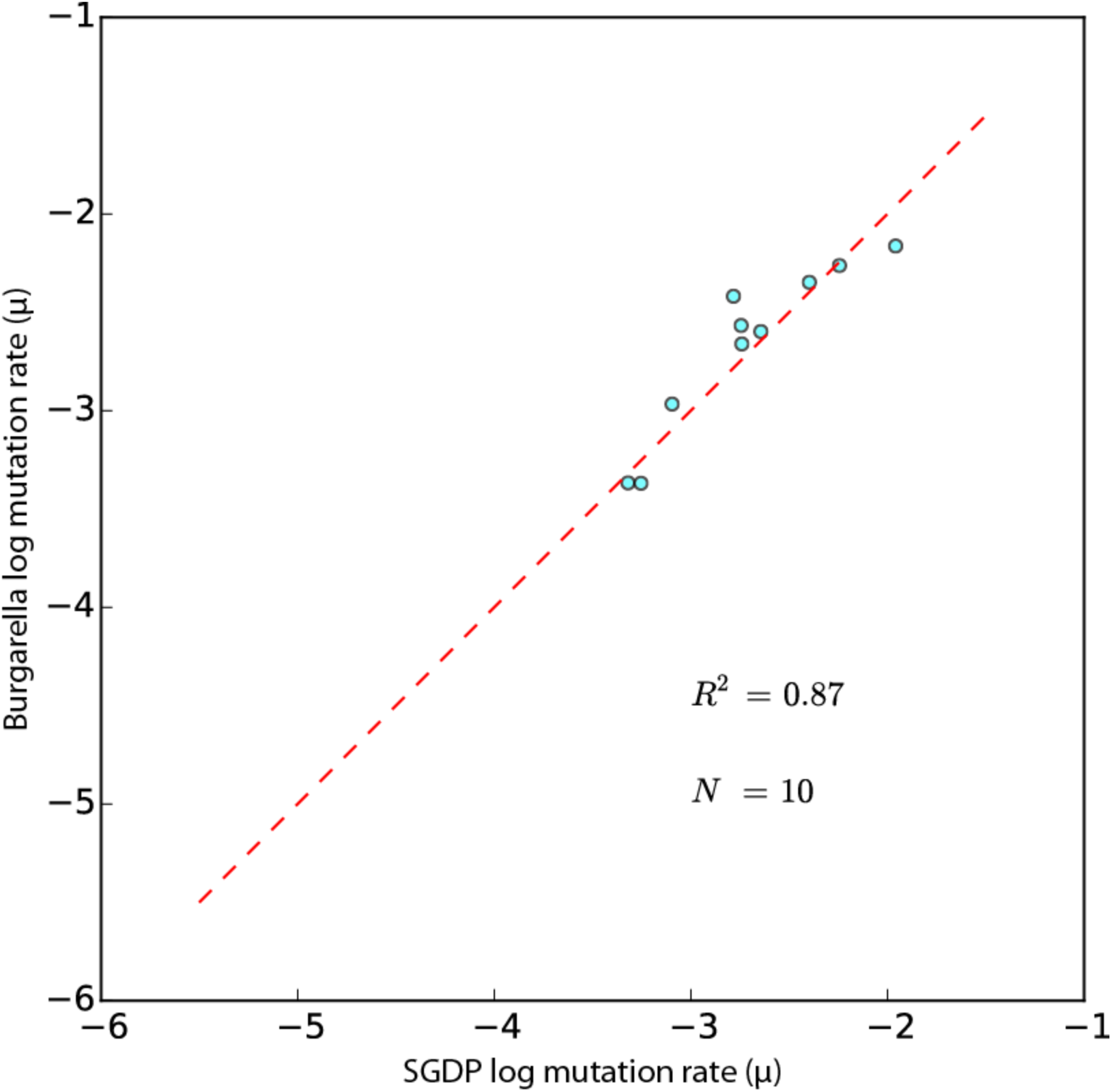
SGDP estimates replicate those Burgarella estimates that were based on large numbers of father-son pairs. The ten mutation rate estimates generated by Burgarella et al. using more than 5,000 father-son pairs are highly concordant with estimates we obtained using the SGDP data.

**Figure S9.**
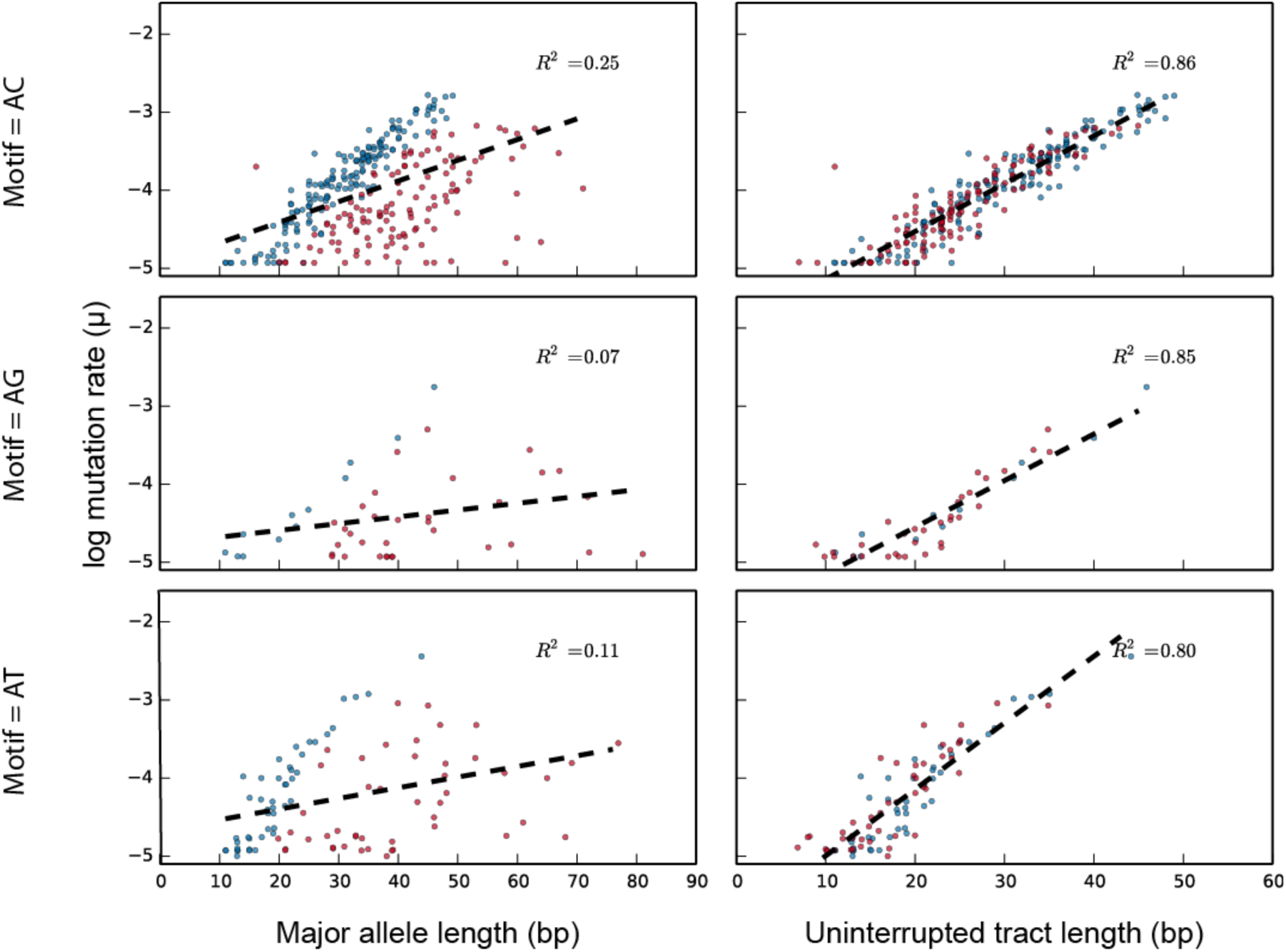
Sequence determinants of Y-STR mutability for loci with dinucleotide repeat units. Stratified by repeat motif (rows), loci with no interruptions to the repeat structure (blue) are generally more mutable than those with one or more interruptions (red). Whereas major allele length is a poor predictor of mutability (first column), the length of the longest uninterrupted tract is a very strong predictor of the log mutation rate for each motif length (second column).

**Figure S10.**
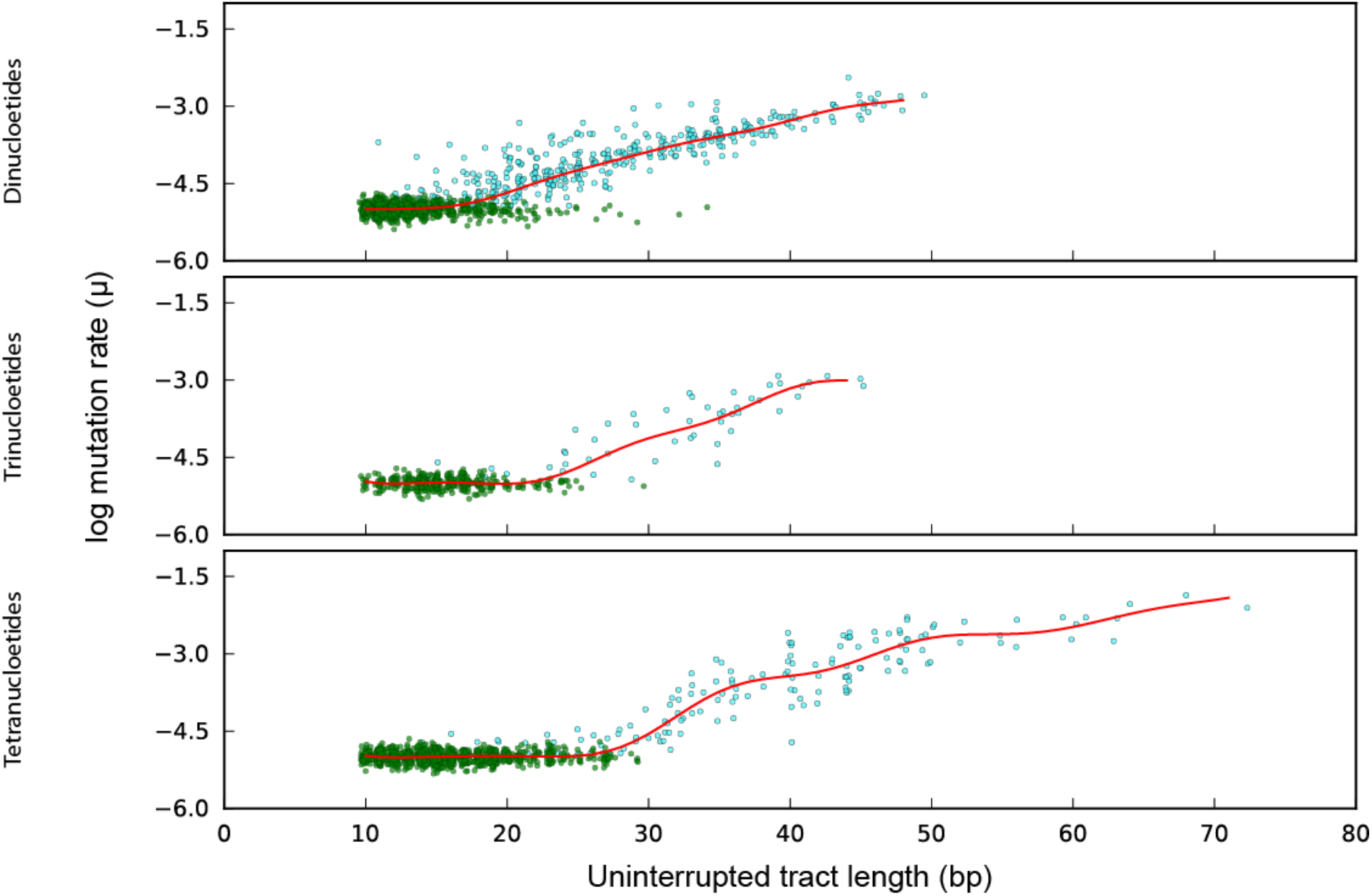
Sequence-based predictors of Y-STR mutation rates. For Y-STRs with di-, tri- and tetranucleotide motifs (rows), the mutation rates for polymorphic Y-STRs (cyan) and fixed Y-STRs (green) were used to fit predictive models of mutation rate (red). In general, the models predict a monotonic increase in log mutation rate with increasing uninterrupted tract lengths. Fixed Y-STRs were assigned a flat rate of 10^−5^ mpg and are displayed using jittered y-values to facilitate visualization.

**Figure S11:**
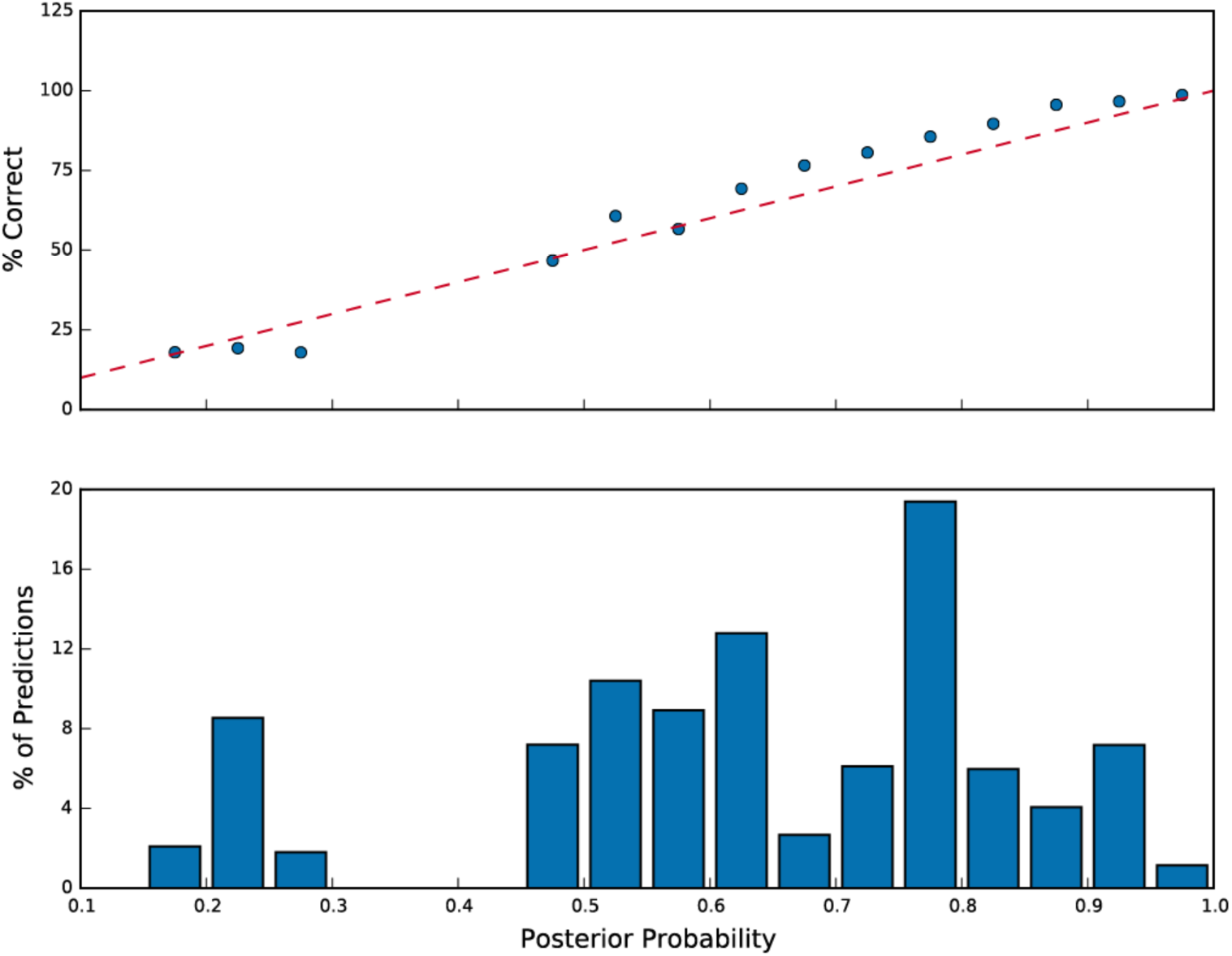
MUTEA-IMPUTE produces well-calibrated imputation probabilities. Y-STR genotypes for 1000 Genomes samples were imputed for loci in the PowerPlex Y23 panel across 1000 iterations, each using a reference panel of 500 samples and an imputation set of 70 samples. The accuracy for each posterior probability bin (upper panel) largely followed the diagonal (red line), demonstrating that the imputation probabilities reflect the true probability of correct imputation.

**Table S3.** Imputation accuracy for loci in the PowerPlex Y23 Panel.

| Marker | $\hat{\mu}$ (mpg) | Posterior > 0% | | Posterior > 70% | |
| --- | --- | --- | --- | --- | --- |
|  |  | % Calls | % Correct | % Calls | % Correct |
| DYS392 | 0.0006 | 100 | 93.1 | 96.4 | 94.5 |
| DYS438 | 0.0006 | 100 | 93.1 | 95.4 | 95.1 |
| DYS437 | 0.0007 | 100 | 92.7 | 95.0 | 94.3 |
| DYS393 | 0.0014 | 100 | 82.5 | 76.9 | 87.8 |
| DYS448 | 0.0017 | 100 | 81.9 | 80.2 | 89.4 |
| DYS533 | 0.0017 | 100 | 78.1 | 74.2 | 85.5 |
| DYS643 | 0.0019 | 100 | 80.0 | 78.1 | 86.3 |
| DYS391 | 0.0022 | 100 | 73.5 | 54.8 | 80.1 |
| Y-GATA-H4 | 0.0023 | 100 | 72.2 | 45.3 | 87.3 |
| DYS390 | 0.0025 | 100 | 76.4 | 51.4 | 84.2 |
| DYS385a | 0.0025 | 100 | 74.3 | 64.5 | 87.9 |
| DYS389I | 0.0027 | 100 | 72.2 | 28.7 | 86.5 |
| DYS19 | 0.0028 | 100 | 70.3 | 35.0 | 88.5 |
| DYS635 | 0.0032 | 100 | 68.6 | 54.9 | 81.7 |
| DYS456 | 0.0037 | 100 | 60.5 | 20.5 | 91.6 |
| DYS549 | 0.0043 | 100 | 55.8 | 4.9 | 85.8 |
| DYS439 | 0.0050 | 100 | 50.0 | 3.0 | 83.5 |
| DYS481 | 0.0051 | 100 | 56.8 | 24.0 | 83.0 |
| DYS385b | 0.0052 | 100 | 53.7 | 17.2 | 87.2 |
| DYS389II | 0.0056 | 100 | 49.1 | 6.6 | 86.8 |
| DYS458 | 0.0079 | 100 | 38.3 | 0.7 | 34.8 |
| DYS570 | 0.0095 | 100 | 41.3 | 0.8 | 94.3 |
| DYS576 | 0.0096 | 100 | 33.6 | 0.5 | 63.9 |
| All |  | 100 | 67.3 | 38.8 | 88.5 |

## References

1. Scally, A., Durbin, R. (2012). Revising the human mutation rate: implications for understanding human evolution. Nature reviews Genetics 13, 745–753.

2. Samocha, K.E., Robinson, E.B., Sanders, S.J., Stevens, C., Sabo, A., McGrath, L.M., Kosmicki, J.A., Rehnstrom, K., Mallick, S., Kirby, A., et al. (2014). A framework for the interpretation of de novo mutation in human disease. Nature genetics 46, 944–950.

3. Kayser, M., and de Knijff, P. (2011). Improving human forensics through advances in genetics, genomics and molecular biology. Nature reviews Genetics 12, 179–192.

4. Conrad, D.F., Keebler, J.E., DePristo, M.A., Lindsay, S.J., Zhang, Y., Casals, F., Idaghdour, Y., Hartl, C.L., Torroja, C., Garimella, K.V., et al. (2011). Variation in genome-wide mutation rates within and between human families. Nature genetics 43, 712–714.

5. Roach, J.C., Glusman, G., Smit, A.F., Huff, C.D., Hubley, R., Shannon, P.T., Rowen, L., Pant, K.P., Goodman, N., Bamshad, M., et al. (2010). Analysis of genetic inheritance in a family quartet by whole-genome sequencing. Science 328, 636–639.

6. Kong, A., Frigge, M.L., Masson, G., Besenbacher, S., Sulem, P., Magnusson, G., Gudjonsson, S.A., Sigurdsson, A., Jonasdottir, A., Jonasdottir, A., et al. (2012). Rate of de novo mutations and the importance of father's age to disease risk. Nature 488, 471–475.

7. Rahbari, R., Wuster, A., Lindsay, S.J., Hardwick, R.J., Alexandrov, L.B., Al Turki, S., Dominiczak, A., Morris, A., Porteous, D., Smith, B., et al. (2016). Timing, rates and spectra of human germline mutation. Nature genetics 48, 126–133.

8. Francioli, L.C., Polak, P.P., Koren, A., Menelaou, A., Chun, S., Renkens, I., Genome of the Netherlands, C., van Duijn, C.M., Swertz, M., Wijmenga, C., et al. (2015). Genome-wide patterns and properties of de novo mutations in humans. Nature genetics 47, 822–826.

9. Itsara, A., Wu, H., Smith, J.D., Nickerson, D.A., Romieu, I., London, S.J., and Eichler, E.E. (2010). De novo rates and selection of large copy number variation. Genome research 20, 1469–1481.

10. Mirkin, S.M. (2007). Expandable DNA repeats and human disease. Nature 447, 932–940.

11. Contente, A., Dittmer, A., Koch, M.C., Roth, J., and Dobbelstein, M. (2002). A polymorphic microsatellite that mediates induction of PIG3 by p53. Nature genetics 30, 315–320.

12. Gebhardt, F., Zanker, K.S., and Brandt, B. (1999). Modulation of epidermal growth factor receptor gene transcription by a polymorphic dinucleotide repeat in intron 1. The Journal of biological chemistry 274, 13176–13180.

13. Shimajiri, S., Arima, N., Tanimoto, A., Murata, Y., Hamada, T., Wang, K.Y., and Sasaguri, Y. (1999). Shortened microsatellite d(CA)21 sequence down-regulates promoter activity of matrix metalloproteinase 9 gene. FEBS letters 455, 70–74.

14. Vinces, M.D., Legendre, M., Caldara, M., Hagihara, M., and Verstrepen, K.J. (2009). Unstable tandem repeats in promoters confer transcriptional evolvability. Science 324, 1213–1216.

15. Sureshkumar, S., Todesco, M., Schneeberger, K., Harilal, R., Balasubramanian, S., and Weigel, D. (2009). A genetic defect caused by a triplet repeat expansion in Arabidopsis thaliana. Science 323, 1060–1063.

16. Weiser, J.N., Love, J.M., and Moxon, E.R. (1989). The molecular mechanism of phase variation of H. influenzae lipopolysaccharide. Cell 59, 657–665.

17. Weber, J.L., and Wong, C. (1993). Mutation of human short tandem repeats. Human molecular genetics 2, 1123–1128.

18. Ellegren, H. (2004). Microsatellites: simple sequences with complex evolution. Nature reviews Genetics 5, 435–445.

19. Sun, J.X., Helgason, A., Masson, G., Ebenesersdottir, S.S., Li, H., Mallick, S., Gnerre, S., Patterson, N., Kong, A., Reich, D., et al. (2012). A direct characterization of human mutation based on microsatellites. Nature genetics 44, 1161–1165.

20. Zhivotovsky, L.A., Underhill, P.A., Cinnioglu, C., Kayser, M., Morar, B., Kivisild, T., Scozzari, R., Cruciani, F., Destro-Bisol, G., Spedini, G., et al. (2004). The effective mutation rate at Y chromosome short tandem repeats, with application to human population-divergence time. American journal of human genetics 74, 50–61.

21. Ballantyne, K.N., Goedbloed, M., Fang, R., Schaap, O., Lao, O., Wollstein, A., Choi, Y., van Duijn, K., Vermeulen, M., Brauer, S., et al. (2010). Mutability of Y-chromosomal microsatellites: rates, characteristics, molecular bases, and forensic implications. American journal of human genetics 87, 341–353.

22. Heyer, E., Puymirat, J., Dieltjes, P., Bakker, E., and de Knijff, P. (1997). Estimating Y chromosome specific microsatellite mutation frequencies using deep rooting pedigrees. Human molecular genetics 6, 799–803.

23. Gymrek, M., Golan, D., Rosset, S., and Erlich, Y. (2012). lobSTR: A short tandem repeat profiler for personal genomes. Genome research 22, 1154–1162.

24. Highnam, G., Franck, C., Martin, A., Stephens, C., Puthige, A., and Mittelman, D. (2013). Accurate human microsatellite genotypes from high-throughput resequencing data using informed error profiles. Nucleic acids research 41, e32.

25. Warshauer, D.H., Lin, D., Hari, K., Jain, R., Davis, C., Larue, B., King, J.L., and Budowle, B. (2013). STRait Razor: a length-based forensic STR allele-calling tool for use with second generation sequencing data. Forensic science international Genetics 7, 409–417.

26. Willems, T., Gymrek, M., Highnam, G., Genomes Project, C., Mittelman, D., and Erlich, Y. (2014). The landscape of human STR variation. Genome research 24, 1894–1904.

27. Genomes Project, C., Auton, A., Brooks, L.D., Durbin, R.M., Garrison, E.P., Kang, H.M., Korbel, J.O., Marchini, J.L., McCarthy, S., McVean, G.A., et al. (2015). A global reference for human genetic variation. Nature 526, 68–74.

28. Sudmant, P.H., Mallick, S., Nelson, B.J., Hormozdiari, F., Krumm, N., Huddleston, J., Coe, B.P., Baker, C., Nordenfelt, S., Bamshad, M., et al. (2015). Global diversity, population stratification, and selection of human copy-number variation. Science 349, aab3761.

29. Gymrek, M. (2016). PCR-free library preparation greatly reduces stutter noise at short tandem repeats. bioRxiv.

30. Danecek, P., Auton, A., Abecasis, G., Albers, C.A., Banks, E., DePristo, M.A., Handsaker, R.E., Lunter, G., Marth, G.T., Sherry, S.T., et al. (2011). The variant call format and VCFtools. Bioinformatics 27, 2156–2158.

31. Stamatakis, A. (2014). RAxML version 8: a tool for phylogenetic analysis and post-analysis of large phylogenies. Bioinformatics 30, 1312–1313.

32. Huson, D.H., Scornavacca, C. (2012). Dendroscope 3: an interactive tool for rooted phylogenetic trees and networks. Systematic biology 61, 1061–1067.

33. Poznik, G.D., Xue, Y., Mendez, F.L., Willems, T.F., Massaia, A., Wilson Sayres, M.A., Ayub, Q., McCarthy, S.A., Narechania, A., and Kashin, S., et al. (2016). Punctuated bursts in human male demography inferred from 1,244 worldwide Y-chromosome sequences. Nature genetics (in press).

34. Helgason, A., Einarsson, A.W., Guethmundsdottir, V.B., Sigurethsson, A., Gunnarsdottir, E.D., Jagadeesan, A., Ebenesersdottir, S.S., Kong, A., and Stefansson, K. (2015). The Y-chromosome point mutation rate in humans. Nature genetics 47, 453–457.

35. Xue, Y., Wang, Q., Long, Q., Ng, B.L., Swerdlow, H., Burton, J., Skuce, C., Taylor, R., Abdellah, Z., Zhao, Y., et al. (2009). Human Y chromosome base-substitution mutation rate measured by direct sequencing in a deep-rooting pedigree. Current biology: CB 19, 1453–1457.

36. Willuweit, S., Roewer, L., and International Forensic, Y.C.U.G. (2007). Y chromosome haplotype reference database (YHRD): update. Forensic science international Genetics 1, 83–87.

37. Benson, G. (1999). Tandem repeats finder: a program to analyze DNA sequences. Nucleic acids research 27, 573–580.

38. Hinrichs, A.S., Karolchik, D., Baertsch, R., Barber, G.P., Bejerano, G., Clawson, H., Diekhans, M., Furey, T.S., Harte, R.A., Hsu, F., et al. (2006). The UCSC Genome Browser Database: update 2006. Nucleic acids research 34, D590–598.

39. Burgarella, C., and Navascues, M. (2011). Mutation rate estimates for 110 Y-chromosome STRs combining population and father-son pair data. European journal of human genetics: EJHG 19, 70–75.

40. Hanson, E.K., and Ballantyne, J. (2006). Comprehensive annotated STR physical map of the human Y chromosome: Forensic implications. Legal medicine 8, 110–120.

41. Li, H. (2013). Aligning sequence reads, clone sequences and assembly contigs with BWA-MEM. arXiv preprint arXiv:13033997.

42. Vermeulen, M., Wollstein, A., van der Gaag, K., Lao, O., Xue, Y., Wang, Q., Roewer, L., Knoblauch, H., Tyler-Smith, C., de Knijff, P., et al. (2009). Improving global and regional resolution of male lineage differentiation by simple single-copy Y-chromosomal short tandem repeat polymorphisms. Forensic science international Genetics 3, 205–213.

43. Purps, J., Siegert, S., Willuweit, S., Nagy, M., Alves, C., Salazar, R., Angustia, S.M., Santos, L.H., Anslinger, K., Bayer, B., et al. (2014). A global analysis of Y-chromosomal haplotype diversity for 23 STR loci. Forensic science international Genetics 12, 12–23.

44. Slatkin, M. (1995). A measure of population subdivision based on microsatellite allele frequencies. Genetics 139, 457–462.

45. Goldstein, D.B., Ruiz Linares, A., Cavalli-Sforza, L.L., and Feldman, M.W. (1995). An evaluation of genetic distances for use with microsatellite loci. Genetics 139, 463–471.

46. Zhivotovsky, L.A., Underhill, P.A., and Feldman, M.W. (2006). Difference between evolutionarily effective and germ line mutation rate due to stochastically varying haplogroup size. Molecular biology and evolution 23, 2268–2270.

47. Felsenstein, J. (1981). Evolutionary trees from DNA sequences: a maximum likelihood approach. Journal of molecular evolution 17, 368–376.

48. Dempster, A.P., Laird, N.M., and Rubin, D.B. (1977). Maximum likelihood from incomplete data via the EM algorithm. Journal of the royal statistical society Series B (methodological), 1–38.

49. Nelder, J.A., and Mead, R. (1965). A Simplex Method for Function Minimization. The Computer Journal 7, 308–313.

50. Brinkmann, B., Klintschar, M., Neuhuber, F., Huhne, J., and Rolf, B. (1998). Mutation rate in human microsatellites: influence of the structure and length of the tandem repeat. American journal of human genetics 62, 1408–1415.

51. Kloosterman, W.P., Francioli, L.C., Hormozdiari, F., Marschall, T., Hehir-Kwa, J.Y., Abdellaoui, A., Lameijer, E.W., Moed, M.H., Koval, V., Renkens, I., et al. (2015). Characteristics of de novo structural changes in the human genome. Genome research 25, 792–801.

52. Ballantyne, K.N., Ralf, A., Aboukhalid, R., Achakzai, N.M., Anjos, M.J., Ayub, Q., Balazic, J., Ballantyne, J., Ballard, D.J., Berger, B., et al. (2014). Toward male individualization with rapidly mutating y-chromosomal short tandem repeats. Human mutation 35, 1021–1032.

53. Nielsen, R. (1997). A likelihood approach to populations samples of microsatellite alleles. Genetics 146, 711–716.

54. Wilson, I.J., and Balding, D.J. (1998). Genealogical inference from microsatellite data. Genetics 150, 499–510.

55. Wilson, I.J., Weale, M.E., and Balding, D.J. (2003). Inferences from DNA data: population histories, evolutionary processes and forensic match probabilities. Journal of the Royal Statistical Society: Series A (Statistics in Society) 166, 155–188 %@ 1467-1985X.

56. Kayser, M., Roewer, L., Hedman, M., Henke, L., Henke, J., Brauer, S., Kruger, C., Krawczak, M., Nagy, M., Dobosz, T., et al. (2000). Characteristics and frequency of germline mutations at microsatellite loci from the human Y chromosome, as revealed by direct observation in father/son pairs. American journal of human genetics 66, 1580–1588.

57. Huang, Q.Y., Xu, F.H., Shen, H., Deng, H.Y., Liu, Y.J., Liu, Y.Z., Li, J.L., Recker, R.R., and Deng, H.W. (2002). Mutation patterns at dinucleotide microsatellite loci in humans. American journal of human genetics 70, 625–634.

58. Xu, X., Peng, M., and Fang, Z. (2000). The direction of microsatellite mutations is dependent upon allele length. Nature genetics 24, 396–399.

59. Ellegren, H. (2000). Heterogeneous mutation processes in human microsatellite DNA sequences. Nature genetics 24, 400–402.

60. Wei, W., Ayub, Q., Xue, Y., and Tyler-Smith, C. (2013). A comparison of Y-chromosomal lineage dating using either resequencing or Y-SNP plus Y-STR genotyping. Forensic science international Genetics 7, 568–572.

61. Kruglyak, S., Durrett, R.T., Schug, M.D., and Aquadro, C.F. (1998). Equilibrium distributions of microsatellite repeat length resulting from a balance between slippage events and point mutations. Proceedings of the National Academy of Sciences of the United States of America 95, 10774–10778.

62. Sainudiin, R., Durrett, R.T., Aquadro, C.F., and Nielsen, R. (2004). Microsatellite mutation models: insights from a comparison of humans and chimpanzees. Genetics 168, 383–395.

63. Petes, T.D., Greenwell, P.W., and Dominska, M. (1997). Stabilization of microsatellite sequences by variant repeats in the yeast Saccharomyces cerevisiae. Genetics 146, 491–498.

64. Bacon, A.L., Farrington, S.M., and Dunlop, M.G. (2000). Sequence interruptions confer differential stability at microsatellite alleles in mismatch repair-deficient cells. Human molecular genetics 9, 2707–2713.

65. Gymrek, M., Willems, T., Guilmatre, A., Zeng, H., Markus, B., Georgiev, S., Daly, M.J., Price, A.L., Pritchard, J.K., Sharp, A.J., et al. (2015). Abundant contribution of short tandem repeats to gene expression variation in humans. Nature genetics.

66. Shao, J., and Wu, C.F.J. (1989). A General Theory for Jackknife Variance Estimation. The Annals of Statistics 17, 1176–1197.

